# Analysis of a Novel Cluster V Mycobacteriophage EniyanLRS and its Therapeutically Relevant LysB

**DOI:** 10.64898/2026.06.26.734815

**Authors:** Kanika Nadar, Kandasamy Eniyan, Urmi Bajpai

## Abstract

Multidrug-resistant tuberculosis and the rising burden of nontuberculous mycobacterial diseases demand innovative treatment strategies. Mycobacteriophages and endolysins offer promising therapeutic potential due to their targeted bacterial cell wall–damaging activity, vast diversity, and lack of antibiotic cross-resistance.

Here, we report the genomic and functional characterisation of a V Cluster mycobacteriophage, ’EniyanLRS’, and its endolysins. Some genomic features: genome size (78.53 kbp), a relatively low GC content (56.9%), a long Tape Measure Protein gene (5.97 kbp), and a high number of tRNAs (24), suggest distinct adaptation and translational efficiency of our phage. Phenotypically, EniyanLRS exhibits a siphovirus morphology, lytic life cycle and infects drug-resistant *Mycobacterium fortuitum*. While its endolysin LysA, harbouring a lysozyme-chitinase-amidase domain, showed moderate lysozyme-like activity without significant antibacterial activity, LysB, an α/β-hydrolase, exhibited superior *in vitro* esterase activity and showed pronounced cell wall disruption in *M. smegmatis* and *M. fortuitum,* along with considerable antibiofilm activity (62.77% and 41.91% inhibition, respectively). Collectively, our analysis reveals key functional properties of EniyanLRS phage and its LysB, demonstrating their suitability as effective biological candidates for mycobacterial therapy.

## Introduction

Mycobacterial cell walls are complex, multi-layered structures that distinguish them from other bacterial species (Lee et al, 1996). The rise in drug-resistant strains of *Mycobacterium tuberculosis* has turned the treatable mycobacterial infection into a persistent crisis, further compounded by limited drug options and high toxicity. The new BPaLM drug regimen offers promising reprieve; however, the associated toxicity and unfavourable outcomes in resistant cases (Ndjeka et al, 2022) remain a concern.

Over the past decade, an increase in the incidence of nontuberculous mycobacteria (NTM) causing both pulmonary and extrapulmonary infections, such as skin and soft tissue infections (SSTI), has been reported (Peeters et al, 2026; Nohreberg et al, 2023; Chung et al, 2018; Honda et al, 2018). In Asia, the examples include Japan, where the prevalence of NTM infection has surpassed that of TB, and South Korea and Taiwan are witnessing a surge in infection rates, especially in NTM pulmonary diseases (Ong et al, 2025). These trends highlight a pressing need to develop novel antimicrobial solutions as a comprehensive public health intervention strategy.

Phages offer significant advantages as antibacterial agents, including their high target specificity, self-replication, and effectiveness against drug-resistant bacterial strains, alone and in combination with standard-of-care antibiotics. In addition, the ability of phages to target biofilms, which can be challenging to eliminate with traditional antibiotics (Zaychikova et al, 2025), is an additional plus. Mycobacteriophages are viruses that infect *Mycobacteria* and are gaining attention as potential therapeutics against drug-resistant mycobacterial infections. The first and the widely reported case of phage therapy for a mycobacterial infection involved a cystic fibrosis patient with a disseminated NTM (*M. abscessus*) infection who was unresponsive to antibiotics. The treatment led to significant clinical improvement, demonstrating the success of mycobacteriophages in treating intracellular bacterial infections (Dedrick et al, 2019). The same group (Dedrick et al, 2023) used mycobacteriophage therapy in 20 patients with NTM infections, where, though mixed results were obtained, the therapeutic phages showed exceptional safety profiles and no evidence of phage resistance, which adds significantly to their promising potential.

Toward the end of the lytic cycle, most mycobacteriophages encode two endolysins: LysA, a peptidoglycan hydrolase, and LysB, a mycolylarabinogalactan esterase that cleaves the bonds between mycolic acid and arabinogalactan (Hatfull, 2010). The enzymes break through the external thick mycolate layer linked with the peptidoglycan layer in mycobacteria to release the phage progeny. The application of mycobacteriophage-encoded endolysins has been successfully evaluated in animal models of human diseases, and their characterisation and development as antimycobacterial candidates are becoming increasingly significant (Abouhmad et al, 2026; Taati et al, 2025; Arora et al, 2024; Singh et al, 2023; Fraga et al, 2019; Park et al, 2018).

Mycobacteriophages are classified into clusters and sub-clusters based on their overall nucleotide sequence similarity (Hatfull, 2018). Among these, Cluster V is one of the least populated, with only 5 members (Table 1). In this study, we report a comprehensive genomic and functional characterisation of ‘EniyanLRS’, a mycobacteriophage that infects *M. smegmatis* and *M. fortuitum*, and its endolysins. The phage genome size is 78.53 kbp, with a GC content of 56.9%, and 148 ORFs. The endolysin sequences, identified from the annotated phage genome, were cloned and purified as recombinant proteins. Their growth-inhibitory effects were assessed on planktonic cells and biofilms of *M. smegmatis* Mc^2^155 and *M. fortuitum* NIHJ1615, and the morphology of the treated cells and biofilms was examined by Field Emission Scanning Electron Microscopy (FESEM).

**Table 1.** General features of V cluster mycobacteriophages (phagesdb.org).

| Phage | Genome (bp) | GC% | 3'Overhang | Number of Genes | tRNA | tmRNA |
| --- | --- | --- | --- | --- | --- | --- |
| EniyanLRS | 78536 | 56.9 | ACCACTGCAAC | 148 | 24 | 1 |
| Cosmo | 78229 | 56.8 | ACCACTGCAAC | 148 | 24 | 1 |
| Azrael100 | 78063 | 56.9 | ACCACTGCAAC | 141 | 21 | 1 |
| MaryV | 76413 | 56.9 | ACCACTGCAAC | 148 | 9 | 1 |
| Wildcat | 78296 | 57.2 | 3' and 11 bases | 148 | 22 | 1 |

## Results and discussion

### Isolation and morphological features of phage EniyanLRS

EniyanLRS mycobacteriophage was isolated from a soil sample collected from Lala Ram Saroop (LRS) TB Hospital (Mehrauli, New Delhi, India) using *M. smegmatis* Mc^2^155 as the primary bacterial host. The plaques were circular and clear, with a thin halo and a diameter of 5.6 mm ± 0.47 (Fig. 1A). TEM analysis revealed a siphovirus morphology; an icosahedral head (58.56 ± 2.45 nm wide) and a 221.515 ± 3.44 nm long non-contractile tail (Fig. 1B). We also observed plaques (Fig. 1C) on a Phx-3-resistant strain of *Mycobacterium fortuitum* (NIHJ1615). Another phage (Wildcat) from Cluster V has also been reported to show a host preference for *M. fortuitum* (Hatfull, 2014). *M. fortuitum* is known to cause cutaneous, pulmonary, post-surgical wound, and corneal infections (Hand et al, 1970). It also causes infections in patients with underlying lung diseases such as TB or bronchiectasis. It has been reported to show resistance to multiple first-line anti-tubercular drugs such as isoniazid, rifampin, and ethambutol, as well as to macrolides and cefoxitin (Shen et al, 2018).

**Figure 1.**
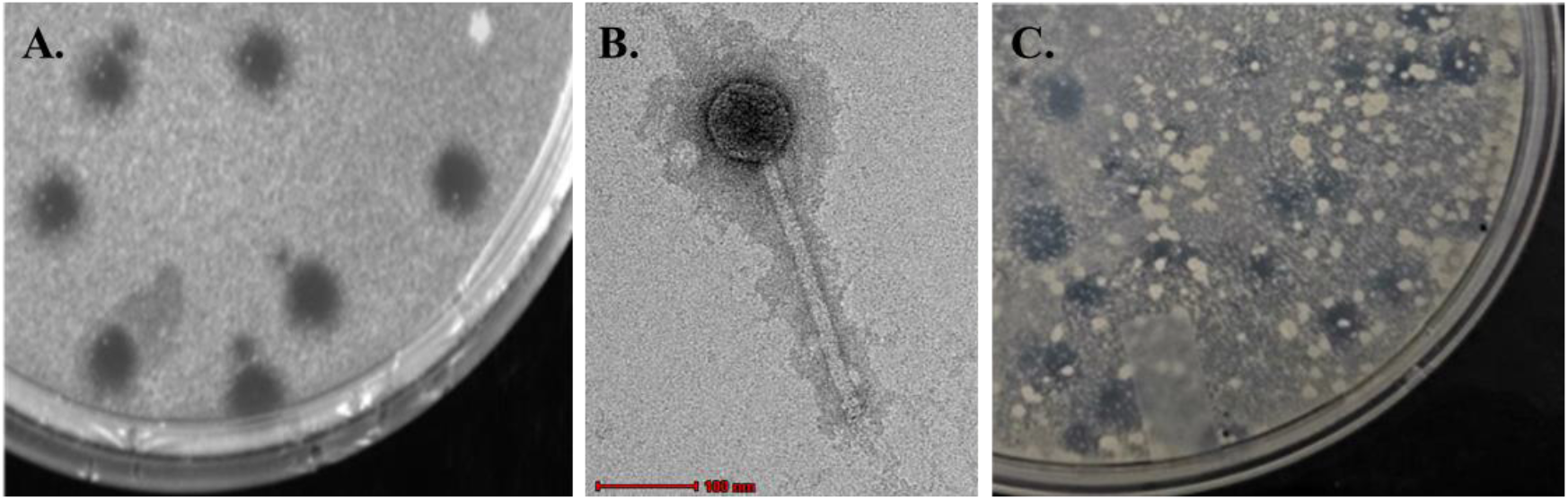
Morphological characterisation of mycobacteriophage EniyanLRS. Representative plate showing **(A)** Plaques (average diameter of 5.6 ± 0.47 mm) on *M. smegmatis* Mc²155 and **(B)** Transmission Electron Micrograph (TEM): Negatively stained with 1% uranyl acetate, the virion head was 58.56 ± 2.45 nm wide, and the tail length was 221.515 ± 3.44 nm. The scale bar represents 100 nm. The morphological measurements were performed using ImageJ software. **(C)** Plaques (average diameter of 4.4 ± 0.52 mm) on *M. fortuitum* NIHJ1615, after 48 h of incubation at 37°C.

### Stability profile

EniyanLRS phage showed viability across a wide range of pH and temperature (Fig. 2). Thermal stability was maintained between 4°C and 55°C, while treatment at 65°C appeared to be a critical threshold, yielding no plaques. EniyanLRS displayed broad pH-range tolerance, with optimal stability between pH 6 and 8 and significant survival at extreme pH values (2, 4, and 10), with titres declining by 18% at pH 2 and 4 and 32% at pH 10.

**Figure 2.**
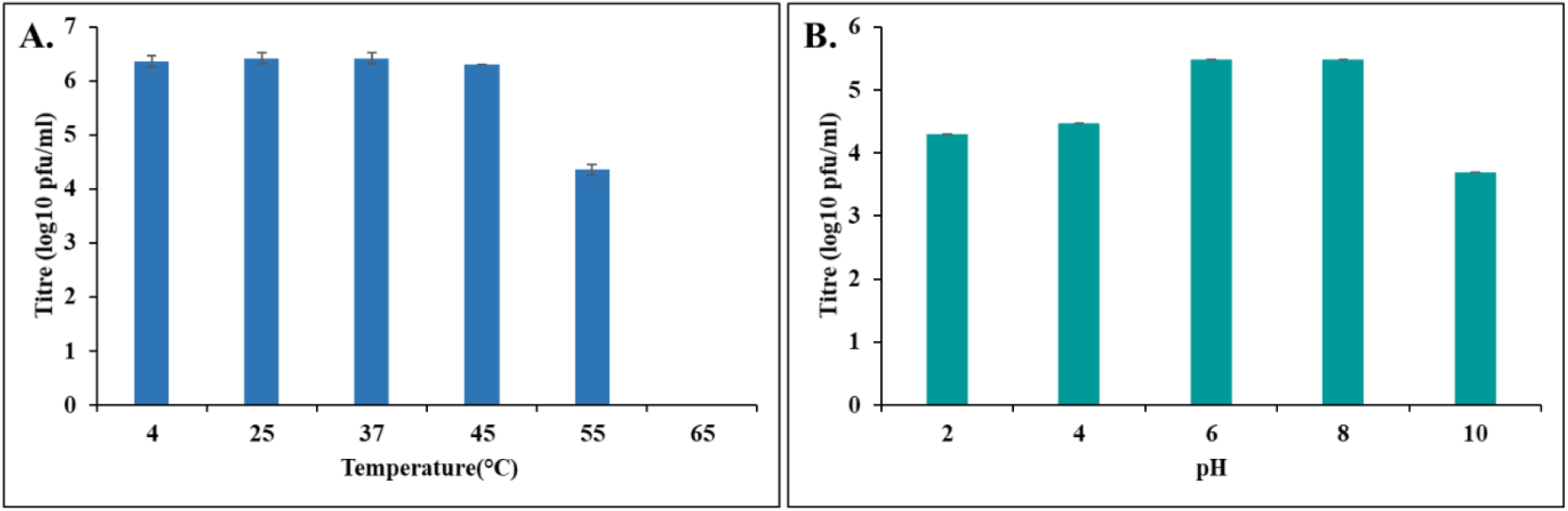
Stability profile of mycobacteriophage EniyanLRS. **(A)** Effect of temperature: Phage viability remained consistent from 4°C to 55°C, with a total loss of infectivity at 65°C. **(B)** Effect of pH: Phage showed optimal stability between pH 6 and 8 and significant survival at pH values 2, 4, and 10. For both assays, phage preparations were incubated under the indicated conditions for 1 h, followed by determining titres on *M. smegmatis* Mc^2^155. Data are expressed as log10 (PFU/mL) and represent the mean of two independent experiments in triplicate (n = 6).

### Phage resistance in bacteria

The emergence of phage resistance in bacteria is one of the major impediments to successful phage therapy. While multi-phage cocktails are considered an effective solution to counter resistance, designing the optimal combination is challenging. Hence, monophage therapy that doesn’t lead to resistance remains a preferred choice.

To determine whether *M. fortuitum* in our study develops resistance to EniyanLRS, we assessed bacterial growth by comparing the OD600 of the infected cultures against uninfected controls. Up to 72 h post-incubation, we noted a significant reduction in OD600 across all tested Multiplicity of Infection (MOIs: 1, 0.1, and 0.01).

At 0.1 and 0.01 MOI, growth inhibition was similar to that observed at MOI 1 for up to 72 h (Fig. 3). However, upon extended incubation (98 h and 122 h), OD600 increased to 0.5 and 0.7 respectively suggesting the emergence of resistance in a subpopulation, although growth remained markedly lower than the uninfected control.

**Figure 3.**
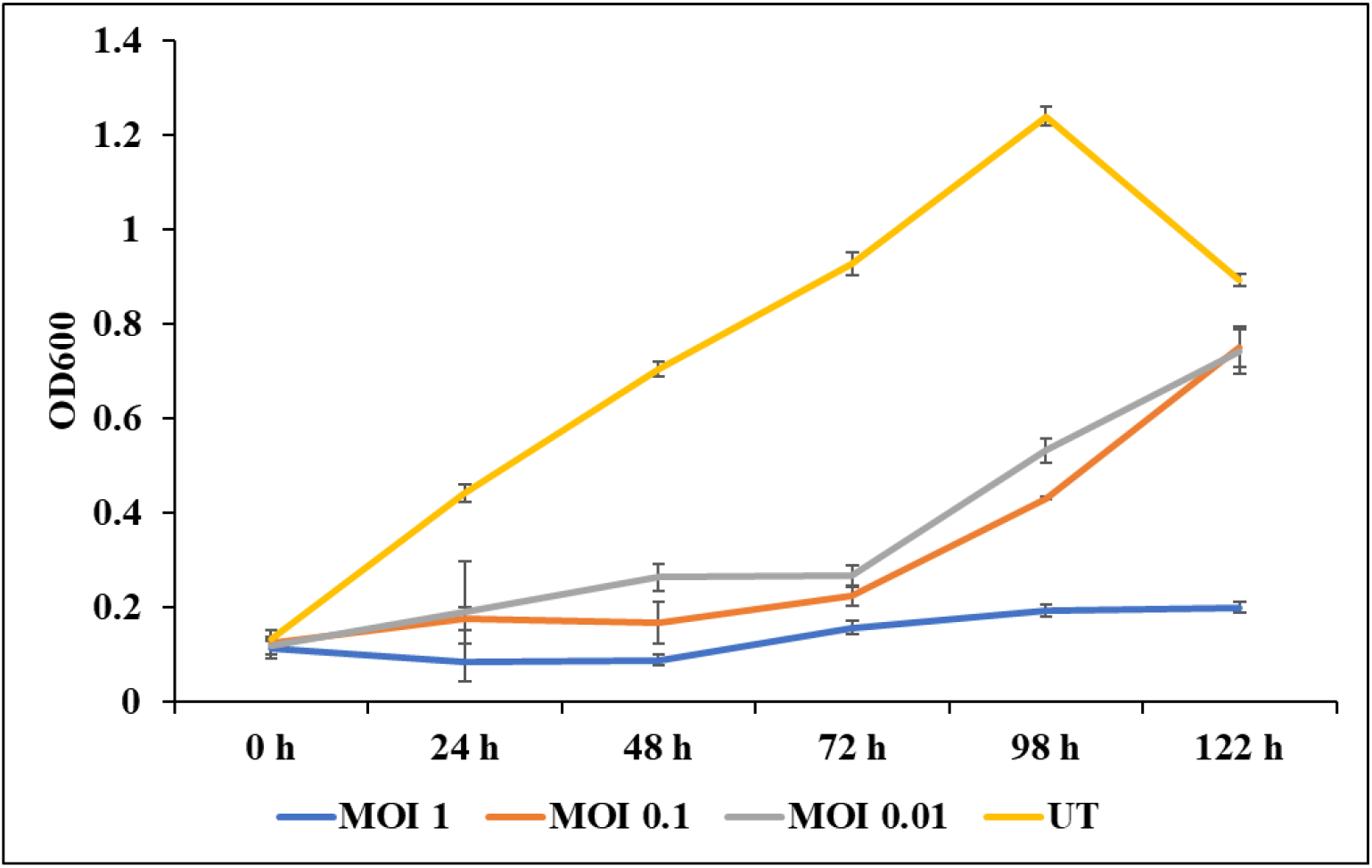
Phage resistance in bacteria. Phage EniyanLRS and *M. fortuitum* were incubated at different MOIs (1, 0.1, 0.01) and OD600 was measured from 0 h to 122 h at an interval of 24 h. At an MOI of 1, bacterial growth remained low throughout whereas at MOI 0.1 and 0.01 the growth started to increase after 72 h. *M. fortuitum* without phage was treated as control. Data is presented as the mean of two independent experiments performed in triplicate (n = 6).

At an MOI of 1, bacterial growth remained low throughout, with only a marginal increase in OD600 from 0.113 to 0.199 at 122 h, indicating sustained inhibition without phage resistance. Concurrently, phage titres increased from an initial 10⁶ PFU/mL to 10^10^ PFU/mL within 24 h, and remained stable thereafter. To corroborate the findings at an MOI of 1, we recovered bacterial cells from phage-treated wells at 122 h and plated them on 7H10 media. Re-exposure of these isolates to the membrane-filtered culture supernatant produced a clear zone of lysis after 48 h at 37°C, confirming susceptibility to EniyanLRS.

These *in vitro* results are encouraging, as low rates of *in vitro* resistance have been suggested to often correlate with *in vivo* efficacy (Dedrick et al, 2021). A subsequent clinical study demonstrated no phage resistance in eleven patients treated with single mycobacteriophage therapy (Dedrick et al, 2023).

### Genome description

The whole-genome sequence of Phage EniyanLRS was characterised using Illumina technology at the Pittsburgh Bacteriophage Institute, University of Pittsburgh, USA. The genome comprises a 78,536 bp double-stranded DNA molecule with a 56.9% GC content, a 11 bp 3’ sticky overhang (ACCACTGCAAC) and 148 predicted open reading frames (ORFs) (Table S1). A significant portion of the genome (58%) consists of hypothetical proteins with unknown functions. These genomic features align closely with those of other Cluster V mycobacteriophages, which typically range from 76 to 78.5 kbp, 141-148 genes, and a GC content between 56.8% and 57.2% (phagesdb.org). A genome map was created using Phamerator tool (https://phamerator.org) (Fig. 4).

**Figure 4.**
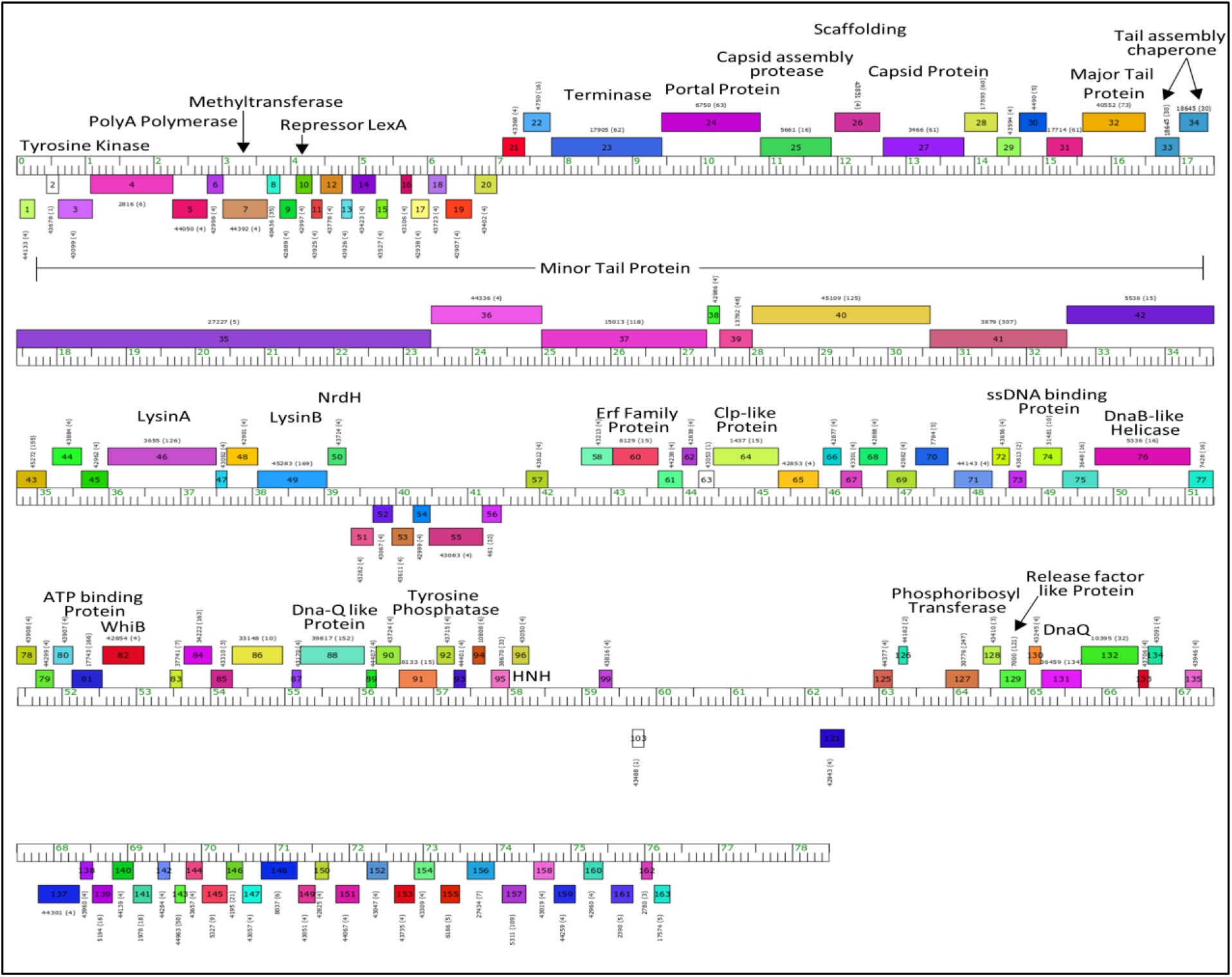
Genome description of mycobacteriophage EniyanLRS. The genome map is generated using the Phamerator tool.

Notably, the EniyanLRS genome encodes 24 tRNAs and one tmRNA. This extensive repertoire suggests that the phage possesses a degree of translational autonomy, potentially enabling it to adjust to host-specific codon biases. This genetic flexibility also likely facilitates its host range. It could be driven by differences in the GC content of the phage (56.9%) compared with that of its mycobacterial hosts: 67.4% and 66% in *M. smegmatis* and *M. fortuitum*, respectively (Machowski et al, 2023) (Supplementary Fig. S1, Supplementary Fig. S2).

To evaluate the biosafety profile and therapeutic suitability of EniyanLRS, we analysed the genome for genes encoding virulence factors, antibiotic resistance, and lysogeny and found the phage to lack virulence factors and antibiotic resistance genes. The absence of repressor and integrase genes (consistent with a lytic lifestyle) further indicated its suitability for therapeutic use.

Cluster V mycobacteriophages represent a unique evolutionary lineage of self-sufficient phages whose genomes diverge significantly from the compact, mosaic architectures of the more populated Clusters (such as A and B). While they retain the canonical siphoviral modular organisation, their larger genomes (≈78–79 kbp) are characterised by a long Tape Measure Protein gene (5.97 kbp), which accounts for their long tails. Unlike the small-genome mycobacteriophages, the lysis cassette in Cluster V phages displays a lateral shift. Furthermore, while a holin gene typically resides within the lysis cassette, its sequence divergence makes it tricky to identify in this Cluster.

Cluster V is currently a sparsely populated group containing only five members. EniyanLRS shares a high identity with the other four members: 99.27% with phages Wildcat and MaryV, 98.96% with Azrael100, and 98.95% with Cosmo. Beyond the V Cluster members, we found the next closest matches to M Cluster phages and unassigned prophages (such as Reindeer at 77.27%, Estes at 74.4%, and prophiGD54-2 at 75.75% identity) (Fig.4A). However, all top hits outside the V Cluster exhibit only 2% query coverage, indicating that the V Cluster phage genomes show significant sequence divergence.

Two of the distinct features of Cluster V phages are their dense tRNA packing (averaging around 20–24 tRNAs) and a low average GC content (56.8% to 57.2%) compared to their hosts, *M. smegmatis* Mc^2^155 (≈67.4%) and *M. fortuitum* NIHJ1615 (≈67.1%). While phages rely on the host’s translation machinery, carrying translation-associated genes, such as tRNAs, confers a fitness advantage. *Mycobacterium* spp. typically prefer GC-ending codons. Conversely, we noted a preference for AT-ending codons, such as GCT, TTA, CTA, CTT, AGA, CGA, CGT, AGT, TCA, TCT, ACT, GTA, and GTT in EniyanLRS. The presence of phage-encoded tRNAs and a tmRNA appears to compensate efficiently for the restricted availability of corresponding host tRNAs. This specialised translation payload reduces reliance on the host’s translational machinery and enables active translation of phage proteins during infection. These adaptations might also confer a broader host range on Cluster V phages, including *M. smegmatis*, *M. fortuitum*, *M. chelonae*, and *M. tuberculosis* (Machowski et al, 2023; Jacobs-Sera et al, 2012; Mageeney et al, 2012; Bailly-Bechet et al, 2007).

Cluster V (and Cluster M) phages exhibit significantly longer TMPs, approximately two-fold larger than those in clusters such as A or F1. Given that the TMP length is directly correlated with tail length, this suggests that Cluster V and M phages possess longer tails. In contrast, phages such as D29 (Cluster A) have shorter TMPs and more compact structural regions, consistent with shorter tail morphology. The genome of Cluster V phages can be distinguished by large numbers of tRNAs and longer TMPs, indicating a coordinated genome architecture that may reflect differences in virion morphology and infection mechanics compared to other clusters (phagesdb.org).

### Phylogenetic analysis and genome comparison

Phylogenetic analysis of the phage genome sequences was performed against a curated database of viral whole-genome sequences across diverse families and host groups using the VipTreeDB Reference Database Version 4.0 (Nishimura et al, 2017). The resulting analysis revealed a close evolutionary relationship between mycobacteriophage EniyanLRS and viruses infecting Actinomycota, with EniyanLRS branching within the same clade as other Cluster V phages (Fig. 5A). This close phylogenetic relationship suggests that these phages possess homologous genetic and functional characteristics, potentially reflecting shared infection and replication mechanisms within *Mycobacterium* hosts. This high degree of conservation was further validated by dot-plot and sequence-alignment analyses, which revealed an exceptionally high sequence identity (95% to 100%) among the phages in Cluster V (Fig. 5B).

**Figure 5.**
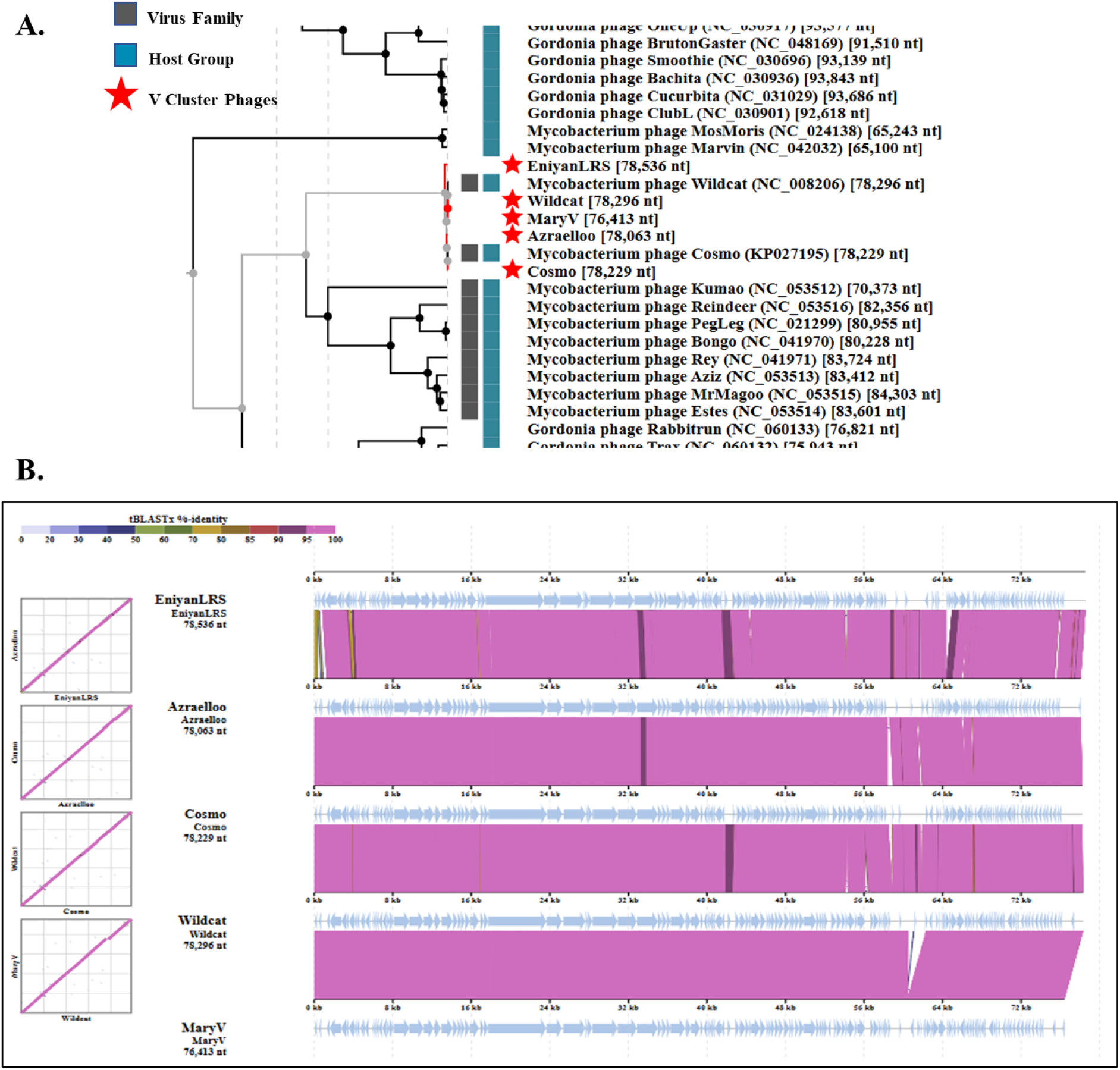
Phylogenetic analysis. **(A)** Proteomic phylogenetic tree generated using ViPTree showing the evolutionary relationship of mycobacteriophage EniyanLRS with related actinobacteriophages based on whole-genome protein similarity. The tree demonstrates clustering of EniyanLRS with other V cluster phages, Wildcat, MaryV, Azrael100, and Cosmo, indicating close evolutionary relatedness. Phages Reindeer and Estes formed comparatively distant branches, suggesting the conservation of a limited core set of proteins despite overall genomic divergence. The blue-coloured bars represent the host group, while the grey coloured bars indicate viral family classification. Red stars denote the selected/reference phages included in the comparative analysis. **(B)** Dot plot and pairwise genome sequence alignment of EniyanLRS with other Cluster V phage genomes (Azrael100, Cosmo, Wildcat, MaryV), highlighting regions of sequence similarity and divergence.

### Lysis cassette

The lysis cassette in EniyanLRS and other Cluster V phages (Fig. 6) displayed a highly conserved arrangement (LysA–HP–Holin–LysB), positioned in the early-to-mid genome (∼31–34% Relative Position Index). TMP is consistently located upstream, indicating modular separation of structural and lysis functions. We also analysed a few mycobacteriophages from different clusters to determine the relative position index (RPI) and organisation of the lysis cassette, which showed clear cluster-dependent variations. The genomic positioning in Clusters F2 (Che9d) and K (TM4, Callalilly) (∼31.5% RPI) was similar to that of cluster V phages, although without a hypothetical protein (LysA–LysB–Holin). Cluster M (Reindeer) showed a relatively earlier placement (∼25% RPI) while retaining the canonical gene order, suggesting conservation with positional shift. More pronounced differences were observed in Clusters A and F1, where the lytic cassette is located in the early genomic region. Cluster A phages (Bxb1, L5, D29) exhibit low RPI values (∼9–12%) with a more compact organisation, ranging from LysA–LysB (A1) to LysA–Holin–LysB (A2). Cluster F1 (Ms6) represents the most extreme case, with the cassette at ∼2% RPI, indicating an unusually early genomic placement. Overall, these comparisons highlight that while the presence of core lysis genes is conserved, their relative positions and gene orders vary across clusters, reflecting evolutionary diversification in genome architecture and potentially distinct lysis-regulation strategies among mycobacteriophages (phagesdb.org).

**Figure 6.**
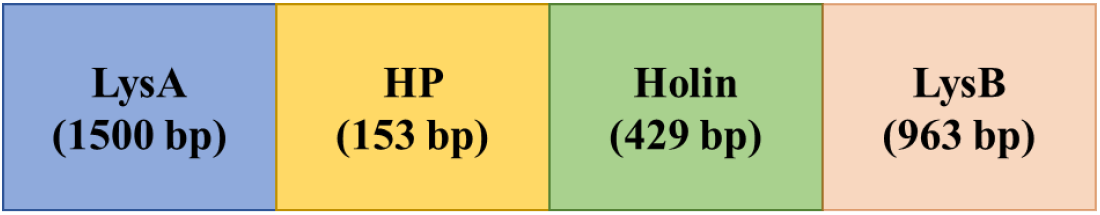
Lysis cassette of mycobacteriophage EniyanLRS. A schematic representation.

### Sequence alignment of EniyanLRS lysin protein sequences

Multiple Sequence Alignment of EniyanLRS LysA with MaryV, Wildcat, Azrael100, and Cosmo showed a high degree of conservation, containing almost entirely identical amino acids with only two conserved and one semi-conserved substitution. In the Cluster A mycobacteriophage Che12, the predicted LysA interaction with the NAG-NAM-NAG trimer within the peptidoglycan glycan chain features a conserved eight-amino acid motif, G-R-[DGT]-X-I-Q-[IL]-[ST], which is found across 19 mycobacteriophage endolysins belonging to Clusters A, B, K and phage Wildcat in Cluster V (Catalao et al, 2018; Saadhali et al, 2016). In the other four Cluster V phages analysed, this conserved region is identified as G-R-G-P-I-Q-L-T (Supplementary Fig. S3). The functional importance and conservation of this motif further confirm its classification within the chitinase protein family (Saadhali et al, 2016). Similarly, multiple sequence alignment of Cluster V phage LysBs demonstrated that the sequences are highly conserved and retain the characteristic GXP motif of cutinases and the pentapeptide G-X-S-X-G motif, both of which are characteristics of serine esterase enzymes (Supplementary Fig. S4). The enzymatic activity of the well-characterised D29 LysB depends on the catalytic triad Ser82−Asp166−His240 (with serine as a constituent of the pentapeptide). Correspondingly, upon evaluating our LysB sequence for these residues, the catalytic triad (Ser-Asp-His) was found to be conserved (Arora et al, 2024).

### Predicted domain organisation of endolysins

EniyanLRS LysA is a 502 amino acid, multi-domain enzyme. Its N-terminal region (residues 16–181) contains a lysozyme domain that functions as an N-acetyl muramidase and embeds a chitinase domain (residues 35–137). This chitinase domain features a GH19 (glycoside hydrolase family 19) motif (residues 69–137), a conserved element across various mycobacteriophages and predominantly found in Cluster A mycobacteriophages, such as Che12 (Catalao et al, 2018). Comparatively, in Che12 LysA, the C-terminal region (residues 215–377) harbours a conserved domain belonging to the peptidoglycan recognition protein (PGRP) superfamily. Within this domain, InterProScan predicts it to contain an amidase catalytic site that cleaves the amide bond between N-acetylmuramic acid and L-alanine residues in the peptidoglycan’s oligopeptide crosslinking chains (Catalao et al, 2011) (Fig. 7A). Conversely, EniyanLRS LysB comprises 321 amino acids and is predicted to contain the characteristic α/β-hydrolase fold (residues 86-318) with an embedded C-linker domain (residues 247-311) (Fig. 7B).

**Figure 7.**
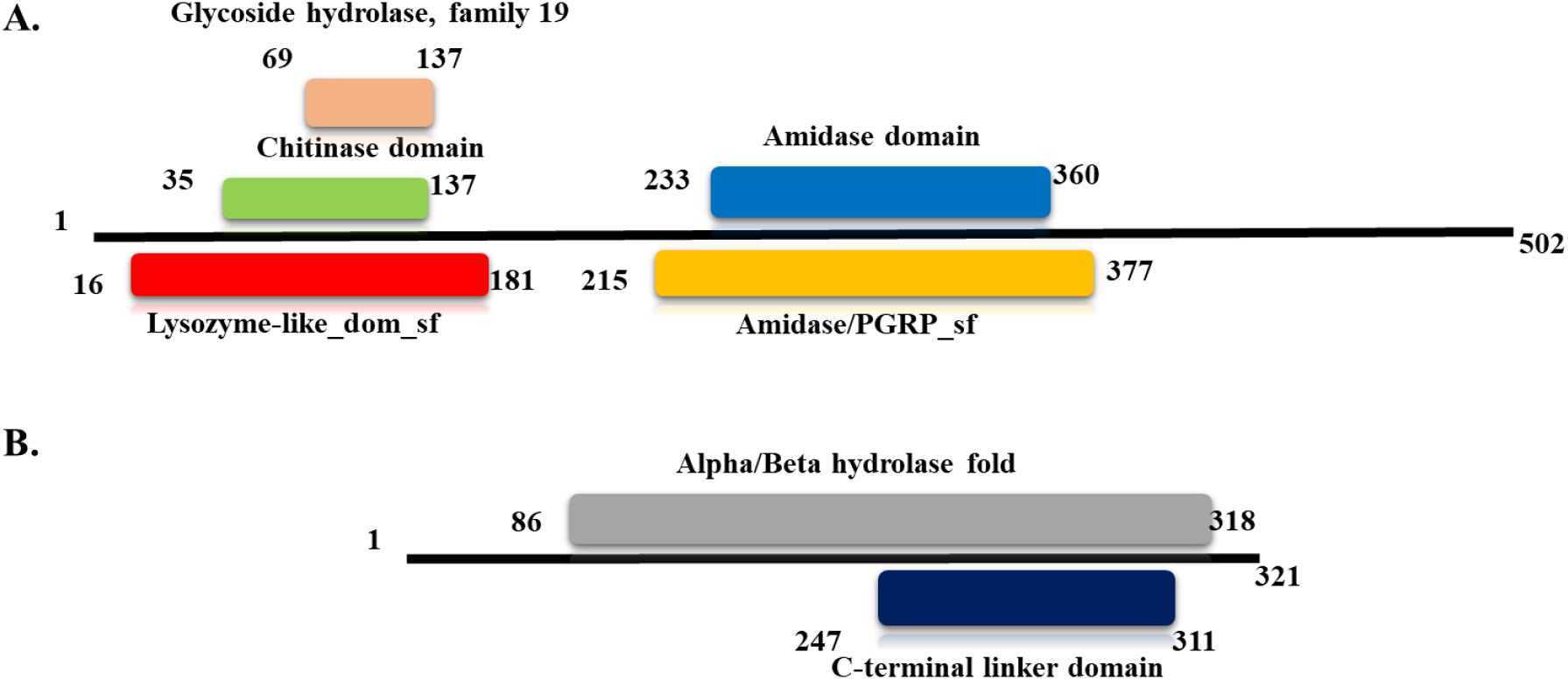
Domain organisation of EniyanLRS endolysins. **(A)** Lysozyme, Chitinase, and Amidase domains in LysA. **(B)** An Alpha/Beta hydrolase fold and a C-linker domain in LysB. The InterProScan tool was used to predict the domains.

### Purification of EniyanLRS lysins and western blot analysis

The His-tagged EniyanLRS LysA and LysB were expressed in *E. coli* BL21 (DE3) and purified by Ni–NTA affinity chromatography (Fig. 8i). Western blotting with anti-His antibodies confirmed the presence of the purified recombinant LysA and LysB (Fig. 8ii).

**Figure 8.**
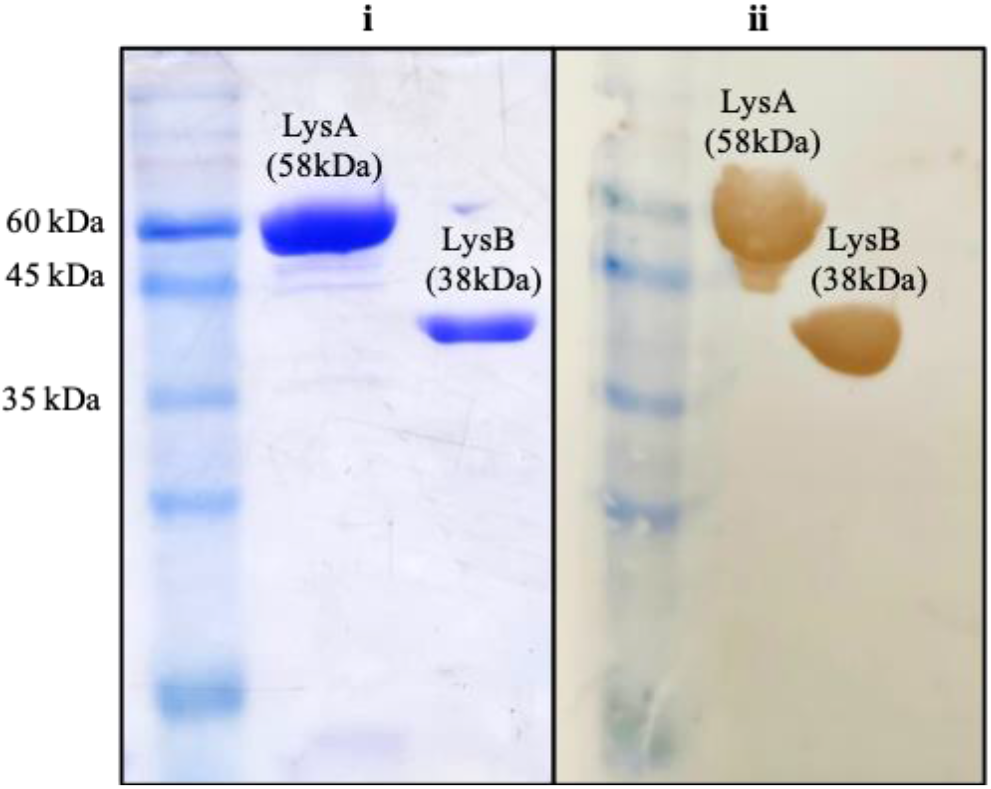
Purification of recombinant endolysins. Analysis of His-tagged Recombinant Endolysins. **(i)** SDS-PAGE. **(ii)** Western Blot.

### Biochemical assays of endolysins

The *in vitro* lysozyme-like activity of EniyanLRS LysA was determined at 37°C using the EnzChek^TM^ Lysozyme Assay kit. Fluorescein-labelled *Micrococcus* cells were used as the substrate, and the relative fluorescence was measured at 485 nm/528 nm (Excitation/Emission). LysB esterase activity was evaluated using 10 mM pNPB. The release of pNP was measured at 410 nm (Fig. 9B) after 30 min of incubation at 37°C.

**Figure 9.**
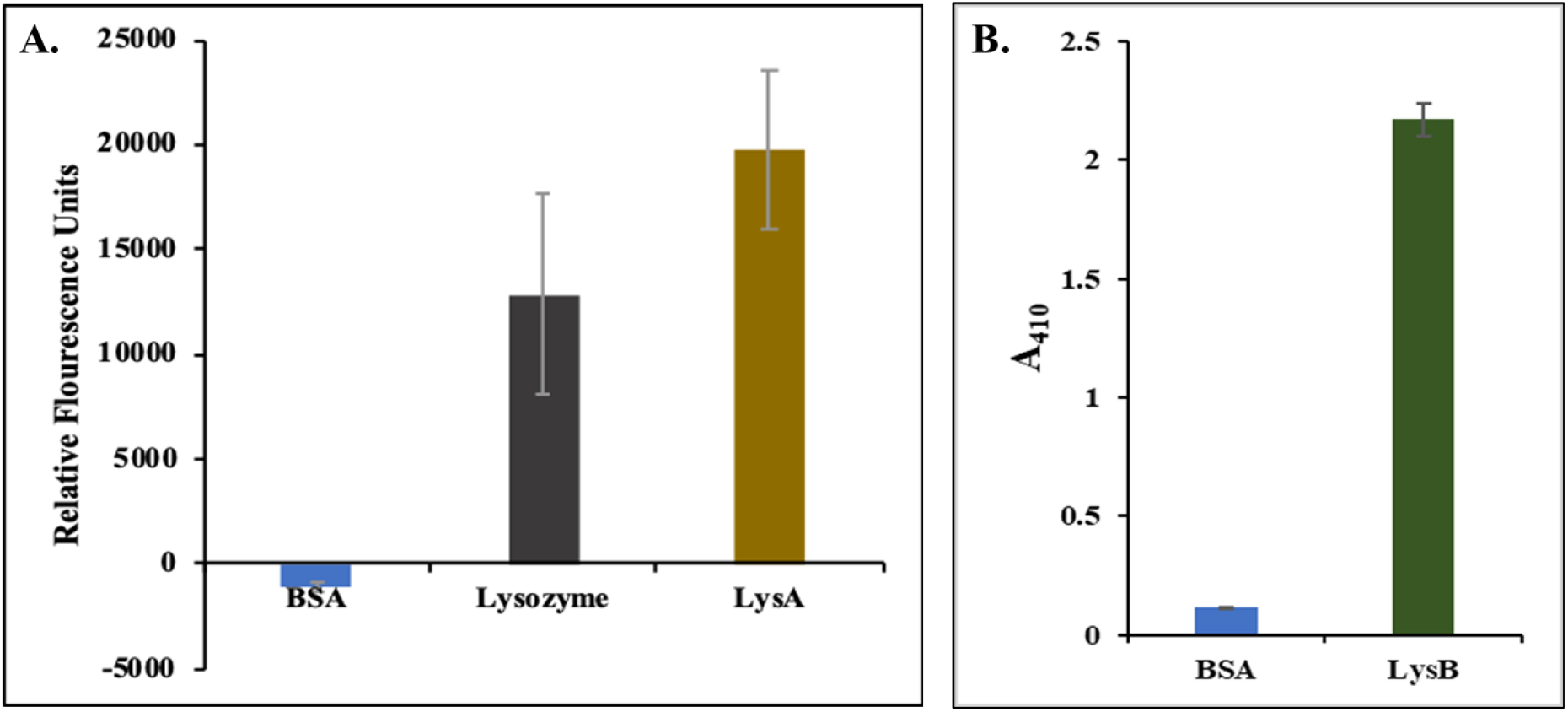
Biochemical assays of EniyanLRS endolysins. **(A)** Lysozyme assay of LysA. Relative Fluorescence Units (RFU) were measured at 485 nm/528 nm (Ex/Em), at a gain of 50. LysA (4 μM); Positive control: Lysozyme (30 U/mL); Negative control: BSA (RFU < 0). The experiment was performed twice independently in triplicate (n = 6). **(B)** Esterase assay of LysB (1 μM) using pNPB (10 mM) was estimated by measuring the release of pNP at 410 nm. Negative control: BSA. The experiments were performed in triplicate (n = 9) three times independently.

Both enzymes demonstrated efficient *in vitro* activity; The lysozyme assay yielded a high relative fluorescence unit (RFU) value for LysA (Fig. 9A), and the specific activity for LysB was calculated to be 5 U/mg.

### Effect of EniyanLRS endolysins against mycobacterial planktonic cells and biofilms

Among the desired properties of native endolysins to be used as therapeutic proteins is their ability to cause lysis on external application (Fischetti et al, 2010). Hence, we evaluated the lytic effect of EniyanLRS LysA and LysB against planktonic cells and biofilm of *M. smegmatis* Mc²155 and a drug-resistant strain (NIHJ1615) of *M. fortuitum*.

#### Planktonic cells

Spot test assays showed a consistent trend across both species: while LysA failed to produce a zone of clearance, LysB yielded distinct lytic zones under identical incubation conditions (48 h at 37°C; Fig. 10A). This confirmed the ability of LysB to damage the mycobacterial cell envelope, likely by cleaving the ester bonds between mycolic acids and arabinogalactan (Payne et al, 2009; Gil et al, 2010).

**Figure 10.**
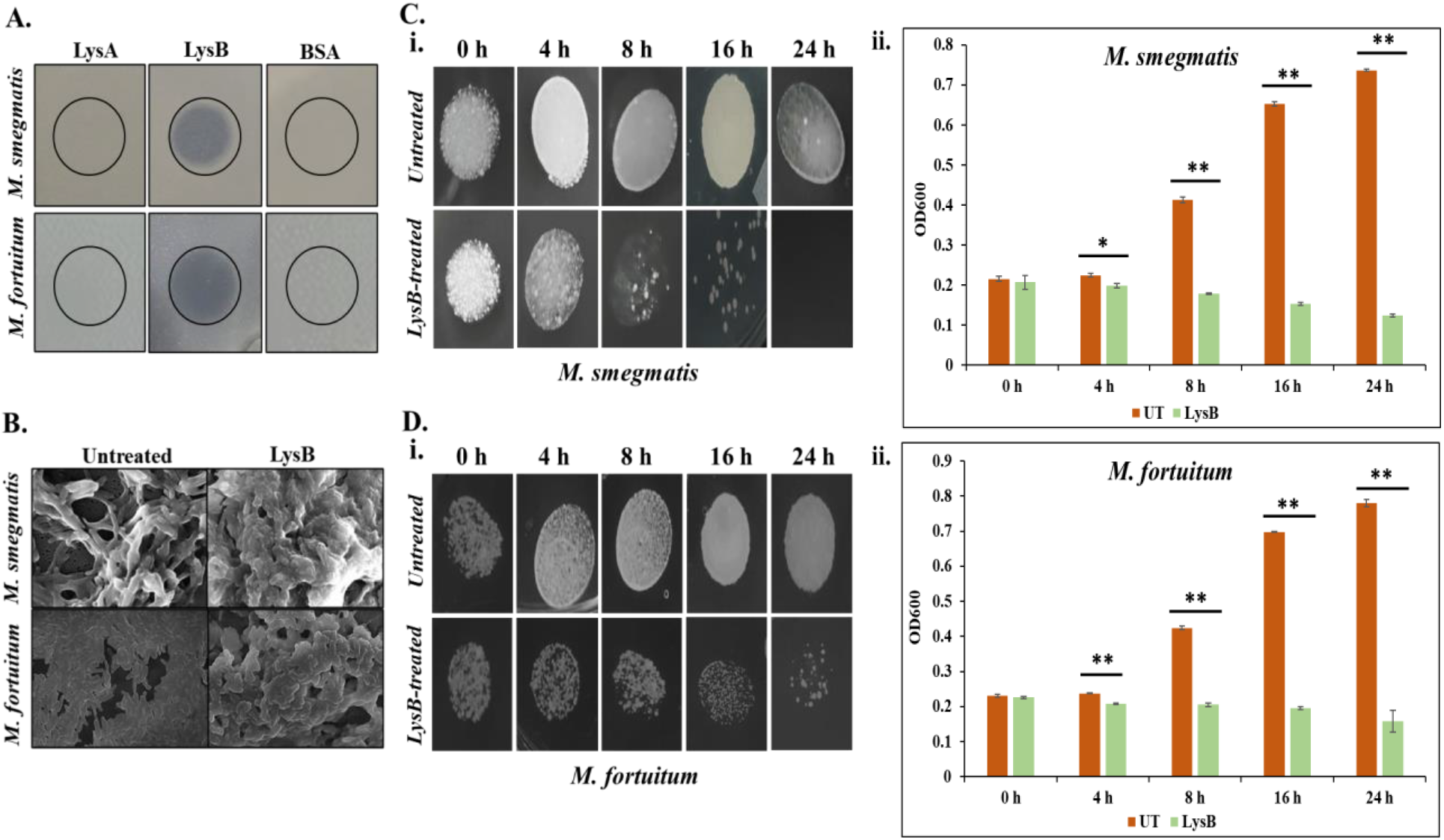
Antibacterial effect of EniyanLRS LysB on planktonic cells of *M. smegmatis* Mc^2^155 and *M. fortuitum* NIHJ1615. **(A)** Spot test showing the antibacterial activity of LysA and LysB. BSA was used as a negative control. **(B)** FESEM images of mycobacterial cells treated with LysB for 4 h at 37°C (15,000 X magnification). **(C)** Time-dependent growth inhibitory effect of LysB on *M. smegmatis*. **(i)** Spot assay: Untreated and LysB-treated cells are depicted at 10^-3^ dilution. **(ii)** TRM measured OD600 at 0 h, 4 h, 16 h and 24 h post-treatment. **(D**) Time-dependent growth inhibitory effect of LysB on *M. fortuitum*. **(i)** Spot assay: Untreated and LysB-treated cells are depicted at 10^-3^ dilution. **(ii)** TRM measured OD600 at 0 h, 4 h, 16 h and 24 h post-treatment. All experiments were performed independently three times in triplicate. Data are presented as mean ± SD (n = 9). Statistical significance between treatment groups was determined by calculating *P* values (** = *P* ˂ 0.01).

#### Turbidity reduction method (TRM)

The lytic activity of LysB was further assessed using a turbidity reduction assay (TRM) at different enzyme concentrations and incubation periods. Based on the concentration-dependent reduction in bacterial growth, a 7.5 µM concentration of LysB was used in subsequent time-dependent lytic activity assays.

LysB-treated *M. smegmatis* cells showed a progressive decrease in OD600 over time, with growth inhibition of 11.5%, 56.7%, 76.4%, and 83.2% observed after 4 h, 8 h, 16 h, and 24 h of incubation, respectively, relative to the untreated control (Fig.9Cii). Similarly, *M. fortuitum* treated with 7.5 µM LysB exhibited growth inhibition of 12.6%, 51.6%, 72.0%, and 79.7% after 4 h, 8 h, 16 h, and 24 h, respectively, compared to the untreated control (Fig. 10Dii). These findings demonstrate a clear time-dependent increase in LysB’s lytic activity against both mycobacterial species. To observe the growth-inhibitory effect of LysB, serial dilutions of LysB-treated *M. smegmatis* and *M. fortuitum* cells were spotted on 7H10 agar plates (Fig. 10Ci, Di). We noted results consistent with the observed trends in turbidity reduction. Furthermore, Field Emission Scanning Electron Microscopy (FESEM) corroborated the antibacterial effect of LysB; compared to the intact morphology of untreated controls, LysB-treated cells of both species showed pronounced surface disruption and distinct cell clumping (Fig. 10B).

#### Biofilm

Biofilms are complex bacterial communities encased in a protective extracellular polymeric substance (EPS) matrix that confer significant resistance to conventional antibiotics (Costerton et al, 1999). We assessed the biofilm inhibitory potential of these enzymes using crystal violet (CV) staining and FESEM imaging. LysA alone expectedly showed no inhibitory effect on biofilm formation in either species. Even in combination with EDTA, a divalent cation chelator, it achieved only a marginal 27.17% inhibition in *M. smegmatis* (Supplementary Fig. S5). In contrast, LysB demonstrated significant antibiofilm activity, independent of EDTA. It achieved a high inhibitory effect of 62.77% against *M. smegmatis* and a lower, yet substantial reduction of 41.91% in *M. fortuitum* biomass (Fig. 11A). FESEM analysis further validated these findings, showing near-complete biofilm inhibition in *M. smegmatis* and moderate reduction in *M. fortuitum* (Fig. 11B).

**Figure 11.**
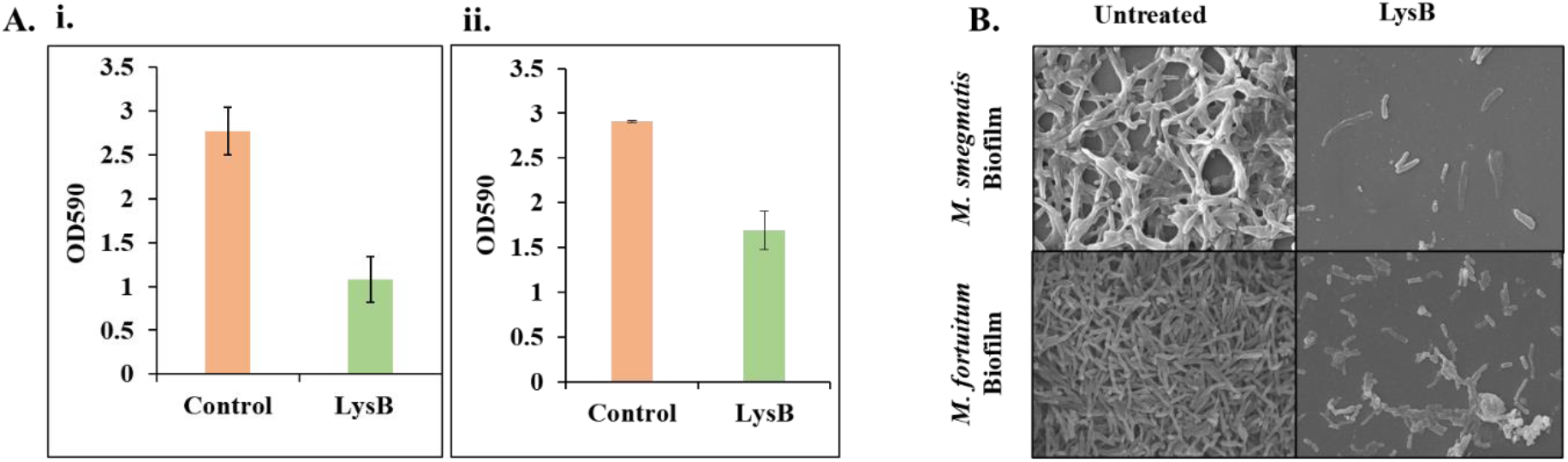
Antibacterial effect of EniyanLRS LysB on biofilm formation in *M. smegmatis* Mc^2^155 and *M. fortuitum* NIHJ1615. **(A)** Crystal Violet Assay measuring the inhibitory effect of LysB (2.5 µM) on biofilm formation **(i)** *M. smegmatis* and **(ii)** *M. fortuitum* after 96 h of incubation at 37°C. Controls represent untreated biofilm formation. Data are presented as the mean of three independent experiments performed in triplicate (n = 9). **(B)** FESEM images (10,000 X magnification) of untreated and LysB (2.5 µM)-treated *M. smegmatis* and *M. fortuitum* biofilms. Incubation was for 96 h at 37°C.

Considering the hydrophobic composition of the mycobacterial cell envelope, it is significant that EniyanLRS LysB not only shows high specific activity compared to other LysBs (Arora et al, 2024), its ability to effectively inhibit growth and biofilm formation in both *M. smegmatis* and *M. fortuitum* further shows its promise as an effective biotherapeutic protein candidate, particularly given the scarcity of literature regarding endolysin activity against mycobacterial biofilms.

## Materials and methods

### Bacterial strains and growth media

*Mycobacterium smegmatis* Mc²155 was used as the host for phage isolation. Cells were grown to mid-log phase in Middlebrook 7H9 broth (HiMedia, India) supplemented with carbenicillin (50 μg/mL), cycloheximide (10 μg/mL), and 0.4% glycerol, with shaking at 37 °C. For solid cultures, Middlebrook 7H10 agar (HiMedia, India) was used. Soil samples were collected aseptically from compost and hospital sites in clean 50 mL centrifuge tubes while wearing gloves. All other chemicals and organic solvents were purchased from HiMedia, India.

### Phage isolation

Soil samples (15–20 g) were mixed with an equal volume of phage buffer, settled for one hour at room temperature, and clarified via centrifugation (2000 × g, 15 min) and 0.45 μm filtration. Phage enrichment was performed by inoculating the filtrate with *M. smegmatis* Mc²155 and 2% LB broth, and incubating overnight at 37°C. The resulting lysate was recovered by centrifugation and 0.22 μm filtration. Phage presence was confirmed via plaque assay on 7H10 agar using a 45-min adsorption period at 37°C. After three rounds of plaque purification, a high-titer lysate was prepared. The purified phage was used to infect a drug-resistant strain (NIHJ1615) of *M. fortuitum*.

### Transmission electron microscopy (TEM)

An aliquot (10 µL) of phage preparation (10^9^-10^10^ PFU/mL) was placed on a carbon-coated copper grid and stained with 2% uranyl acetate. Phage morphology was observed using a JEOL TEM-1400 (JEOL Ltd.) at 120 kV, and images were taken at a magnification of 30,000X at SAIF, AIIMS, New Delhi. The phage morphological measurements were performed using ImageJ software.

### Stability analysis

To examine the effect of temperature on phage viability, the phage lysate (3 x 10^6^ PFU/mL) was incubated at 4°C, 25°C, 37°C, 45°C, 55°C, 65°C and 75°C for 1 h. To study the effect of pH, the phage preparation was diluted into Tris buffer adjusted to pH 2, 4, 6, 8, and 10, and incubated for 1 h at 37°C. Treated phages were diluted and plated on 7H10. Post-incubation, the phage titre was determined using standard plaque assays. The experiments were performed twice independently in triplicate (n = 6).

### Phage resistance assay

To study phage resistance in *M. fortuitum*, a time-dependent assay for quantitative measurement of bacterial growth at OD600 was performed. Briefly, *M. fortuitum* culture, grown to mid-log phase and diluted to 0.1 OD, was infected with EniyanLRS at different Multiplicity of Infection (MOIs) of 1, 0.1, and 0.01 in microtiter plates. Incubation was at 37°C under shaking conditions (200 rpm) for a 122-h period. Every 24 h, OD600 was measured, and an aliquot was drawn to enumerate PFU/mL. The experiments were performed in triplicate and repeated independently twice (n = 6). The results are calculated as the mean ± SD relative to the uninfected control.

To verify whether surviving bacteria acquired resistance, we recovered cells from phage-treated wells after 122 h of incubation and isolated them on Middlebrook 7H10 agar plates. Concurrently, the culture supernatant from the 122 h wells was harvested and membrane-filtered (0.22 μm) to collect free phage particles. The bacterial culture was plated onto fresh 7H10 agar to form a bacterial lawn, and dilutions of the filtered supernatant were spotted onto the lawn and incubated at 37°C for 48 h to inspect for zones of lysis.

### Whole genome sequencing (WGS) and annotation

For WGS, phage genomic DNA was isolated from a high-titer lysate (10^10^–10^11^ PFU/mL) using a modified Phenol-Chloroform-Isoamyl alcohol (PCI) protocol (phagesdb.org) as described by Arora et al. (2025). Concentration and DNA purity were validated via A260/A280 and A260/A230 ratios using a multimode plate reader (Synergy LX, BioTek).

WGS of the phage was performed at Prof Graham F Hatfull’s Pittsburgh Bacteriophage Institute’s sequencing facility using the Illumina platform. Raw sequence quality was assessed using FastQC. The genome sequence was assembled using Newbler and Consed, and automatic annotation was performed with DNA Master v. 5.0.2, followed by manual curation. The final assembled genome was obtained as FASTA files and used for subsequent annotation. DNA Master (https://phagesdb.org/DNAMaster/) was used to predict the CDS region, and functional annotation was performed using HMMER (Finn et al, 2011) and BLAST. All programs were run using default parameters. The sequenced and annotated genome was submitted to GenBank (https://www.ncbi.nlm.nih.gov/genbank/) under accession KY385381.1, named ‘EniyanLRS’. The genome was analysed for acquired antimicrobial resistance genes using ResFinder 4.7.2 (https://genepi.food.dtu.dk/resfinder); PhageLeads (https://phageleads.dk/) and PhageScope (https://phagescope.deepomics.org/) were used to predict phage lifestyle and virulence factors.

### Cloning and expression of endolysins

LysA and LysB genes were PCR-amplified using EniyanLRS phage gDNA as the template. Genes were cloned into the pET28a vector (Novagen, USA) and expressed in *E. coli* BL21 (DE3) as described in our previous study (Eniyan et al. 2020). Cultures were grown in Luria Broth to an OD600 of 0.4–0.6 and induced with 0.1 mM IPTG at 16°C overnight (200 rpm). Post-induction, cells were harvested and resuspended in lysis buffer (25 mM Tris, pH 7.9, 0.5 M NaCl, 10 mM imidazole, 1 mM PMSF). Lysis was performed by sonication (90% amplitude, ten 30-second on/off cycles), followed by centrifugation at 5000 × g for 45 min at 4°C to separate the soluble and pellet fractions.

Recombinant His-tagged endolysins were purified from the soluble fraction using Ni-NTA affinity chromatography. The lysate was incubated with Ni-NTA agarose for 2 h at 4°C, packed into a column, and washed with 20 mM and 40 mM imidazole. Target proteins were eluted with 250 mM imidazole, analysed by 12% SDS-PAGE, and dialysed into 25 mM Tris and 150 mM NaCl. For Western blot validation, purified proteins were transferred to a PVDF membrane at 90 V for 2 h. The membrane was blocked with 3% BSA and probed with a mouse anti-6x His primary antibody (Santa Cruz Biotech, USA) at 1:2000 for 4 h, followed by an HRP-conjugated anti-mouse IgG secondary antibody (1:10,000) for 45 min. Following standardised PBS and PBS-T (0.05% Tween-80) washes, the blot was developed using 3,3-diaminobenzidine (DAB) and H2O2 (Eniyan et al, 2020; Mahmood & Yang, 2012).

### Endolysin activity

#### Biochemical assays

The lysozyme assay for LysA was performed according to the manufacturer’s protocol (EnzChek Lysozyme Assay Kit). Briefly, 50 µL of the substrate (50 µg/mL) was incubated with LysA enzyme (4 µM) in a total reaction volume of 100 µL for 1 h at 37°C. After incubation, fluorescence was measured in a plate reader (Synergy LX, BioTek) at 485 nm/528 nm (Ex/Em). Lysozyme (30 U/mL) was taken as a positive control, and BSA (50 µg) as a negative control. Each experiment was performed twice independently in triplicate (n = 6), and the results were calculated as mean ± standard deviation (SD).

The esterase activity of the LysB enzyme was assessed using the p-nitrophenol butyrate (pNPB) assay, as described by Fraga et al. (2019) with minor modifications. Release of p-nitrophenol (PNP) was measured by incubating purified LysB (1 µM) with 10 mM pNPB in 25 mM Tris buffer (pH 7.2) in a total reaction volume of 200 µl for 30 min at 37°C, followed by measuring the OD at 410 nm. BSA (1 mg/mL) was used as a blank. The specific activity (U/mg) was calculated using the PNP standard plot. Each experiment was performed in triplicate and repeated independently three times (n = 9), and the results were calculated as mean ± SD.

#### Antibacterial activity using the spot test

Endolysins (10 µg each of LysA & LysB) were spotted on *M. smegmatis* Mc^2^155 lawn on 7H10 plates containing CB (50 μg/mL), CHX (10 μg/mL), and glycerol (0.4%) and *M. fortuitum* NIHJ1615 plates containing CHX (10 μg/mL) and glycerol (0.4%). The plates were incubated at 37°C for about 48 h to observe the formation of a clearance zone. For LysB, Tween-80 (0.05%) was added to the media.

#### Turbidity reduction method (TRM)

TRM studies were conducted as discussed in Arora et al. (2024). The effect of LysB was checked on *M. smegmatis* Mc^2^155 and *M. fortuitum* NIHJ1615 cells. Briefly, the cells were adjusted to an OD600 of 0.3 and treated with different concentrations of LysB (2.5 μM, 5 μM, 7.5 μM and 10 μM) for a fixed time duration of 24 h. The incubation was done at 37°C with shaking at 200 rpm. Untreated cells served as the control. The treated and the untreated cells were serially diluted and spotted on 7H10 plates containing Tween-80 (0.05%) and incubated at 37°C for 48 h. For the time-interval experiments, a fixed concentration (7.5 μM) of LysB was incubated for different time intervals (4 h, 8 h, 16 h, and 24 h). Each experiment was performed twice independently in triplicate (n = 6).

#### FESEM analysis of LysB-treated *M. smegmatis* Mc^2^155 and *M. fortuitum* NIHJ1615 cells

Cells grown in Tween-80 containing Middlebrook 7H9 complete medium were treated with LysB (7.5 μM) for 4 h at 37°C. After incubation, treated cells were centrifuged at 11,000 × g for 5 min, followed by a single wash with phosphate buffer saline (PBS). Washed cells were fixed overnight at 4°C in 2.5% glutaraldehyde and dehydrated using an ethanol gradient (5 min each in 60%, 80%, and 90%, and 10 min in 100%). The dehydrated cell pellet was resuspended in Tween-containing PBS and was inoculated on a 0.2 µm polycarbonate (PC) membrane. The membrane was air-dried and mounted on a stub for gold sputter coating. Samples were visualised at 15,000× magnification using a TESCAN (at the Indian Institute of Technology, New Delhi) (Lai et al, 2015).

#### Biofilm-inhibitory effect of EniyanLRS lysins

*M. smegmatis* Mc^2^155 and *M. fortuitum* NIHJ1615 biofilms were formed in 24-well plates following the protocol of Das et al. (2024). To study the inhibitory effect of LysA (with and without 3 mM EDTA) on biofilm formation, cells (OD600 = 0.1) were mixed with lysin (2.5 µM) in microtiter plate wells and incubated for 96 h at 37°C under static conditions. For LysB, the effect was studied with Tween-80 (0.05%) and without EDTA. Biofilms were quantified using the Crystal Violet (CV) staining method (Kiefer et al, 2015).

#### Crystal violet (CV) staining

After the Lysin treatment, the unattached cells were aspirated, the biofilms were air-dried and stained with freshly prepared CV solution (1%). After 30 min, the CV was aspirated, and the wells were air-dried, washed twice with 1X PBS and air-dried again. The CV stain was extracted with 95% ethanol for 20 min at 37°C, and the absorbance was measured at 590 nm (Arora et al, 2024). The experiments were performed in triplicate, repeated independently three times (n = 9), and the results were calculated as the average percentage inhibition ± SD relative to the untreated biofilm.

#### FESEM analysis of *M. smegmatis* Mc^2^155 and *M. fortuitum* NIHJ1615 biofilm

LysB-treated biofilm was grown on 12 mm coverslips in a 24-well plate for 96 h at 37°C, followed by fixation overnight at 4°C in 2.5% glutaraldehyde (Siddam et al, 2021). Fixed biofilm was washed with PBS and then dehydrated using a gradient of ethanol (once for 10 min each at 30%, 50%, 70%, 85%, and 95%, and thrice in 100%). Dehydrated biofilm was washed with PBS and treated with a 1:1 Hexamethyldisilazane (HMDS)/ethanol mixture for 10 min, followed by 100% HMDS for 1 h. The coverslip was air-dried and mounted on a stub for gold sputter coating, and the biofilm was visualised at 10,000X magnification using a TESCAN (at the Indian Institute of Technology, New Delhi) to analyse biofilm inhibition or any morphological damage to the biofilm-forming cells. Untreated biofilm served as the control.

## Conclusion

This report communicates the first study on comprehensive characterisation of a Cluster V mycobacteriophage and presents evidence for the therapeutic potential of its LysB enzyme. The isolation of EniyanLRS from a clinical hospital environment highlights the role ecological diversity can play in isolating phages against pathogenic species of *Mycobacterium*.

Genomic screening confirmed that the EniyanLRS genome is devoid of virulence factors, antibiotic resistance genes, and lysogeny-related marker genes, including integrase and att sites. These findings, corroborated by the observation of clear plaques, suggest a lytic lifestyle and establish that the phage can be used for therapeutic applications. Our analysis suggests that EniyanLRS and other Cluster V phages exhibit high structural complexity due to their large genome sizes. While higher RPI and long TMP genes are characteristic features of large-genome mycobacteriophages, the large pool of tRNA genes serves as an evolutionary adaptation to compensate for translational constraints arising from the GC-content divergence between the phage and its host. By compensating for these significant codon usage biases, Cluster V phage can overcome the host constraints.

Functional dissection of the phage-encoded endolysins revealed a clear functional divergence between LysA and LysB. While LysA possesses a highly conserved chitinase-amidase architecture and demonstrable enzymatic activity *in vitro*, its limited ability to lyse intact mycobacterial cells highlights the formidable permeability barrier imposed by the mycomembrane. In contrast, LysB emerged as an effective antimycobacterial enzyme capable of acting “from without,” and disrupting the mycolylarabinogalactan linkage critical for cell wall integrity. Its superior esterase activity, broad host efficacy, and pronounced antibiofilm action, with visible structural damage, collectively distinguish LysB as a promising enzybiotic candidate. Its activity on both planktonic cells and biofilm in *M. smegmatis* and the pathogenic NTM *M. fortuitum* highlights its potential in mycobacterial disease management, where biofilm-associated persistence and intrinsic drug resistance limit therapeutic success.

Phylogenetic analysis revealed that LysB proteins from Cluster V mycobacteriophages, including the EniyanLRS LysB, cluster closely with LysinB_MF, a recently reported prophage-encoded LysB from our laboratory (Das et al, 2025). Both of these LysBs have been experimentally observed to exhibit lytic activity against *M. fortuitum*. Although these proteins originate from different genomic contexts (the prophage that encodes LysinB_MF is a truncated 9.4 kb fragment and remains unassigned to a Cluster), their close relationship is likely a consequence of host-specific selective pressure acting on LysB evolution. LysB enzymes hydrolyse the ester linkage between mycolic acids and the arabinogalactan layer of the mycobacterial cell envelope, and efficient lysis requires enzymes to adapt to the specific cell wall structure of each host. The presence of a closely related LysB in a prophage and a Cluster-assigned phage suggests that adaptation to the *M. fortuitum* cell envelope may constrain sequence divergence in this enzyme. This observation supports the idea that host specialisation, rather than extensive horizontal gene transfer, plays a key role in shaping lysis modules in Cluster V mycobacteriophages. Such host-driven conservation may also contribute to the limited diversity observed within this Cluster.

In conclusion, this study not only expands the understanding of a novel V-Cluster mycobacteriophage, but also reinforces the growing acceptance that phage-derived lysins can target complex, drug-resistant pathogens. Overall, this work positions EniyanLRS phage and its LysB as prominent candidates for future translational development and contributes to next-generation antimycobacterial therapeutic strategies.

## Supplementary Information

**Table S1.**
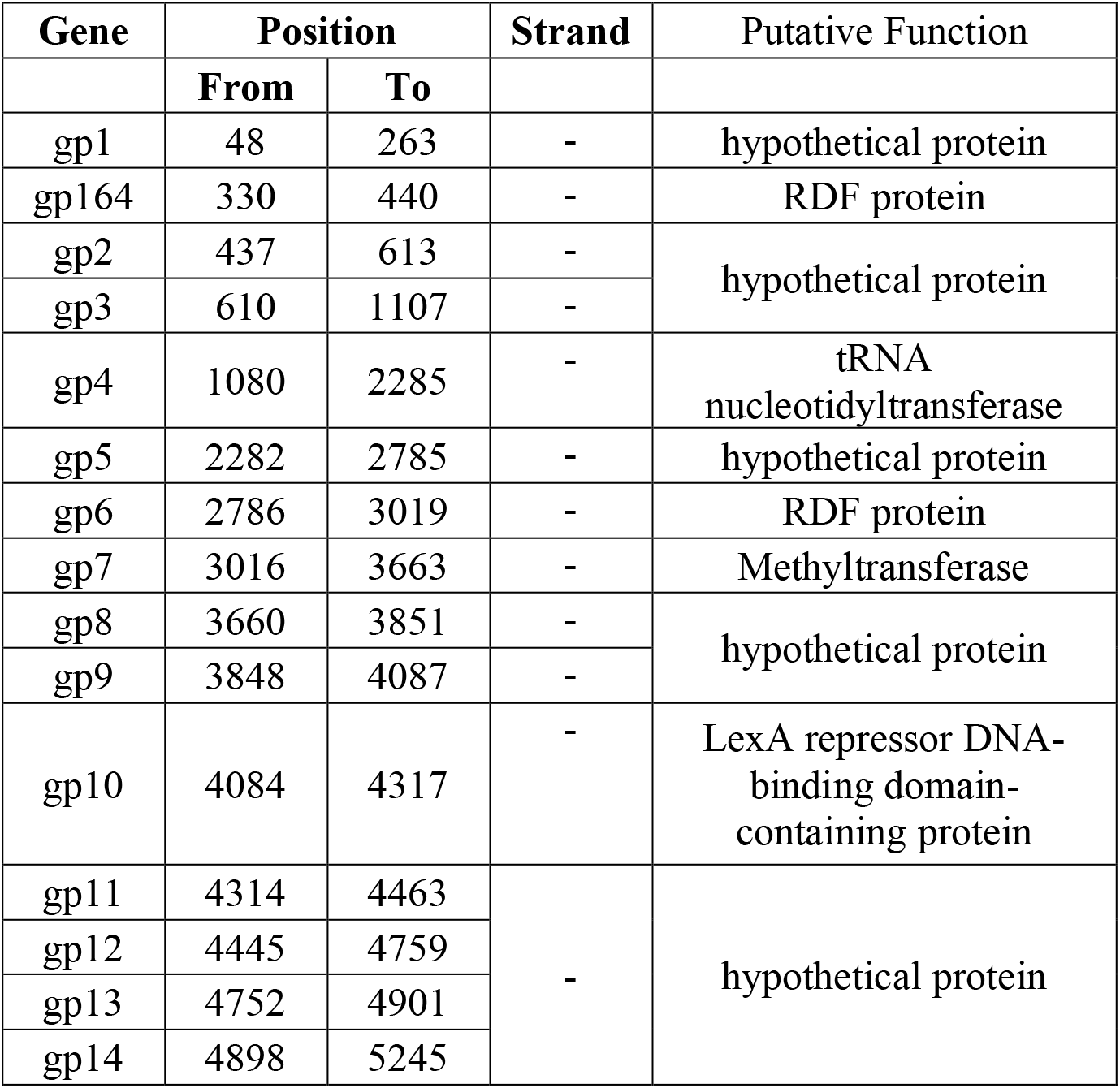
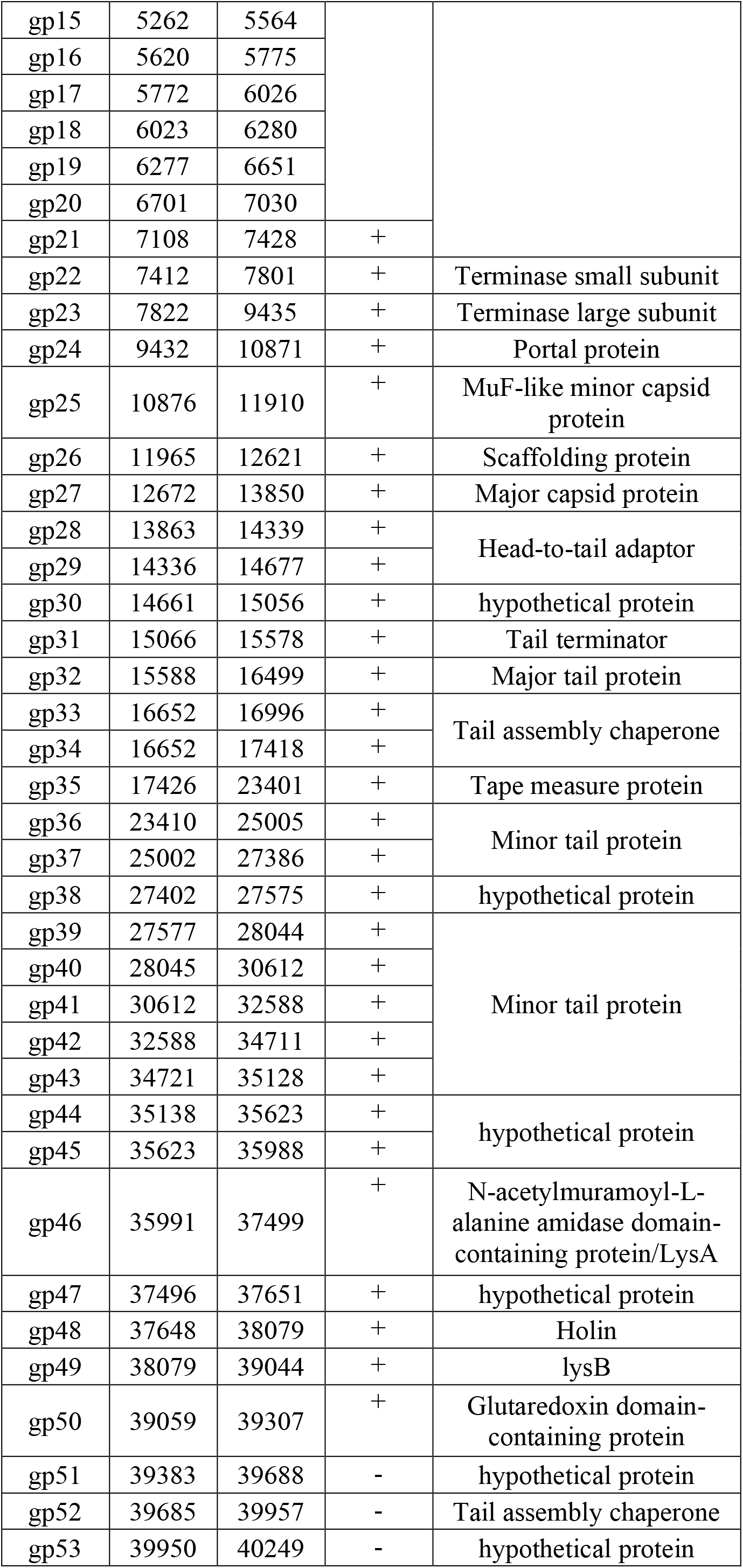
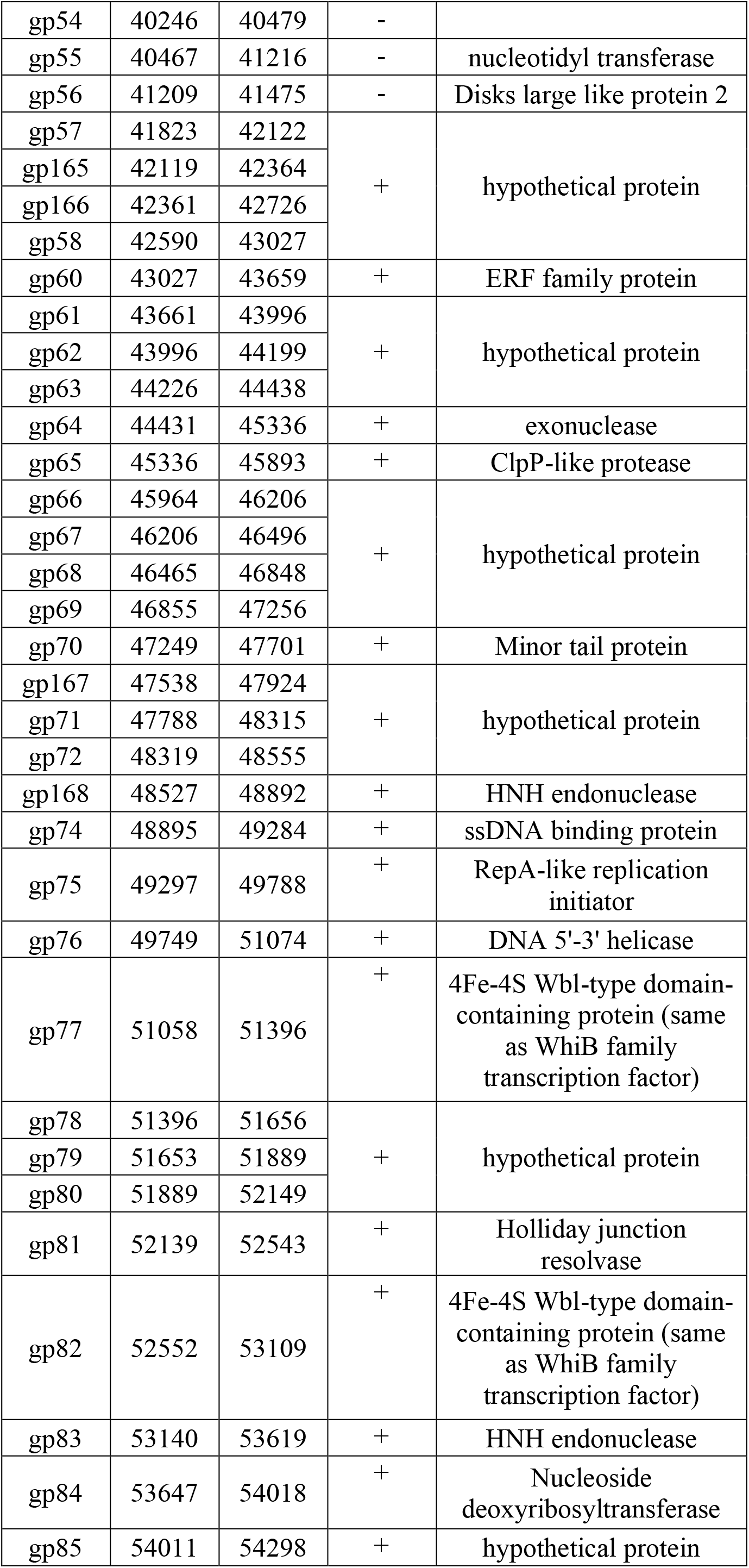
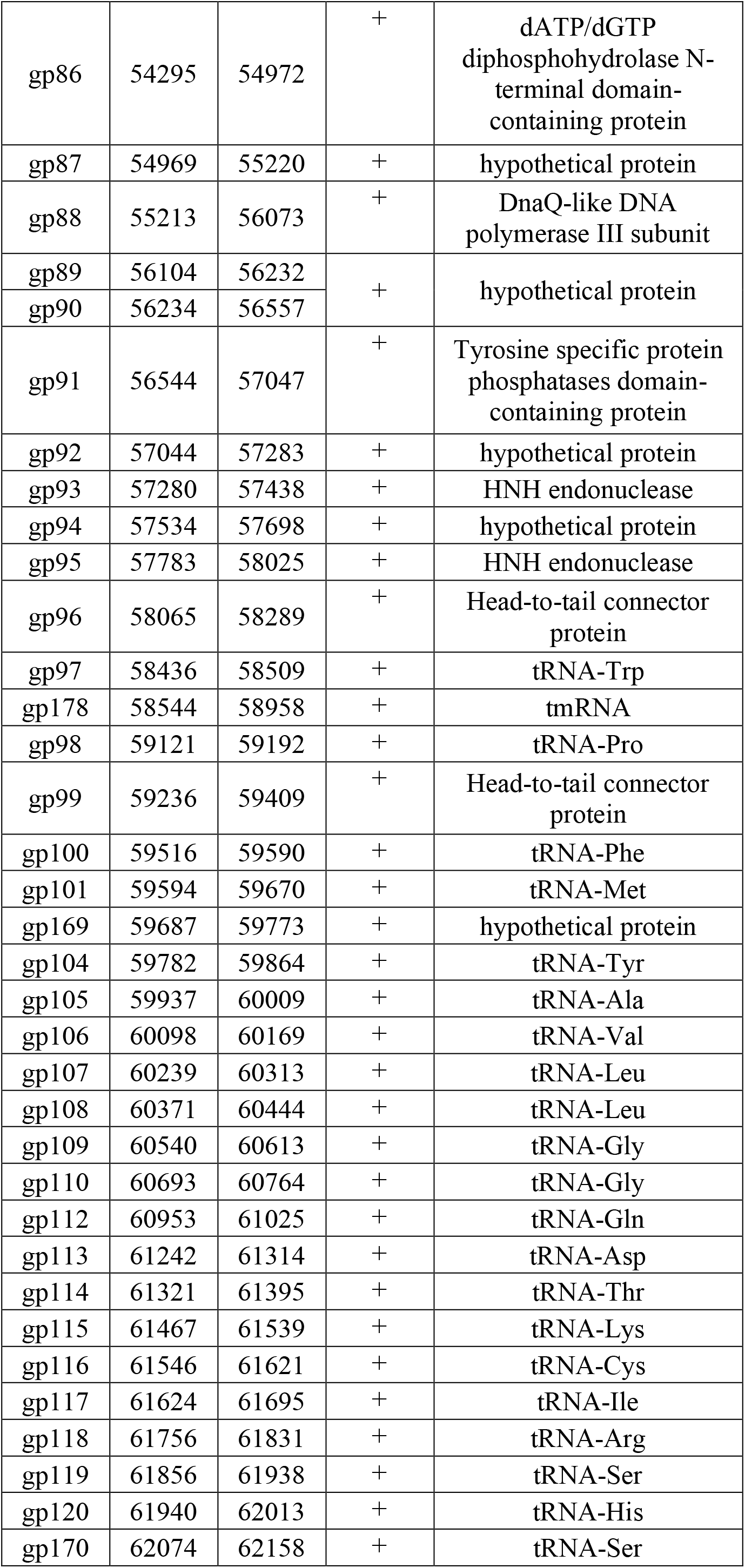
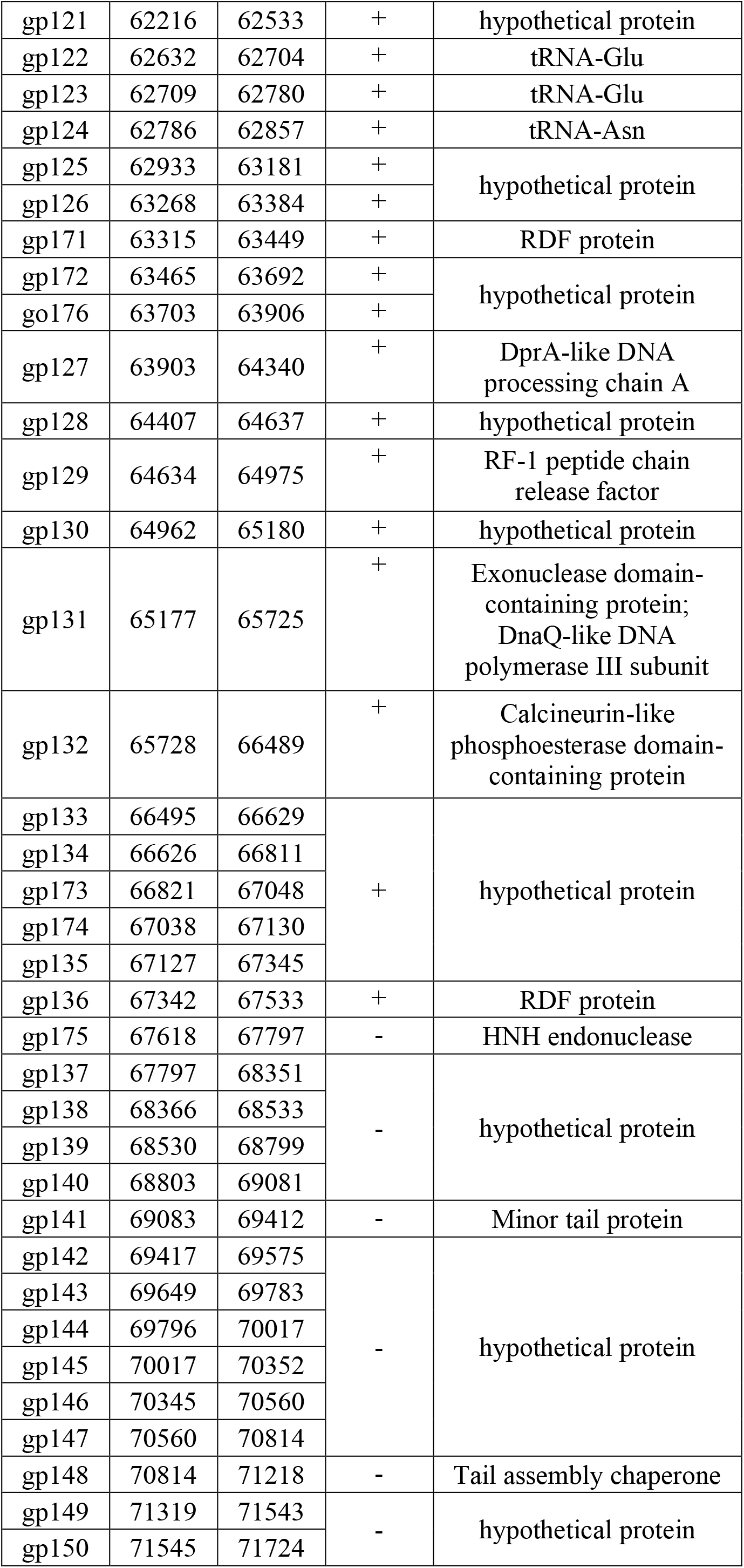
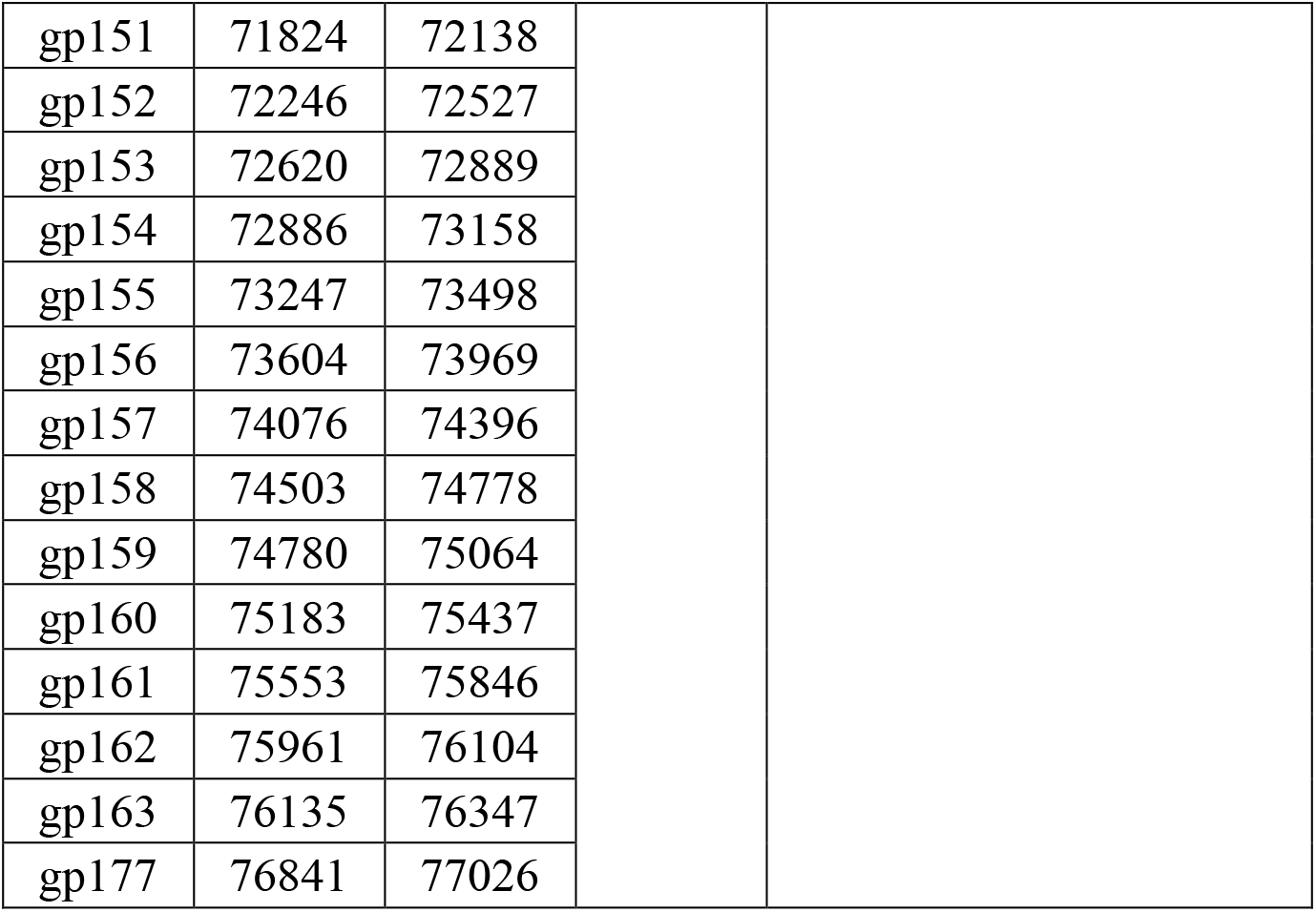
Functional annotation of Mycobacteriophage EniyanLRS Genome. This table lists the predicted ORFs of mycobacteriophage EniyanLRS along with their genomic positions, strand orientation and putative functions. HMMER tool (https://www.ebi.ac.uk/Tools/hmmer/home) was used for functional annotation.

**Figure S1.**
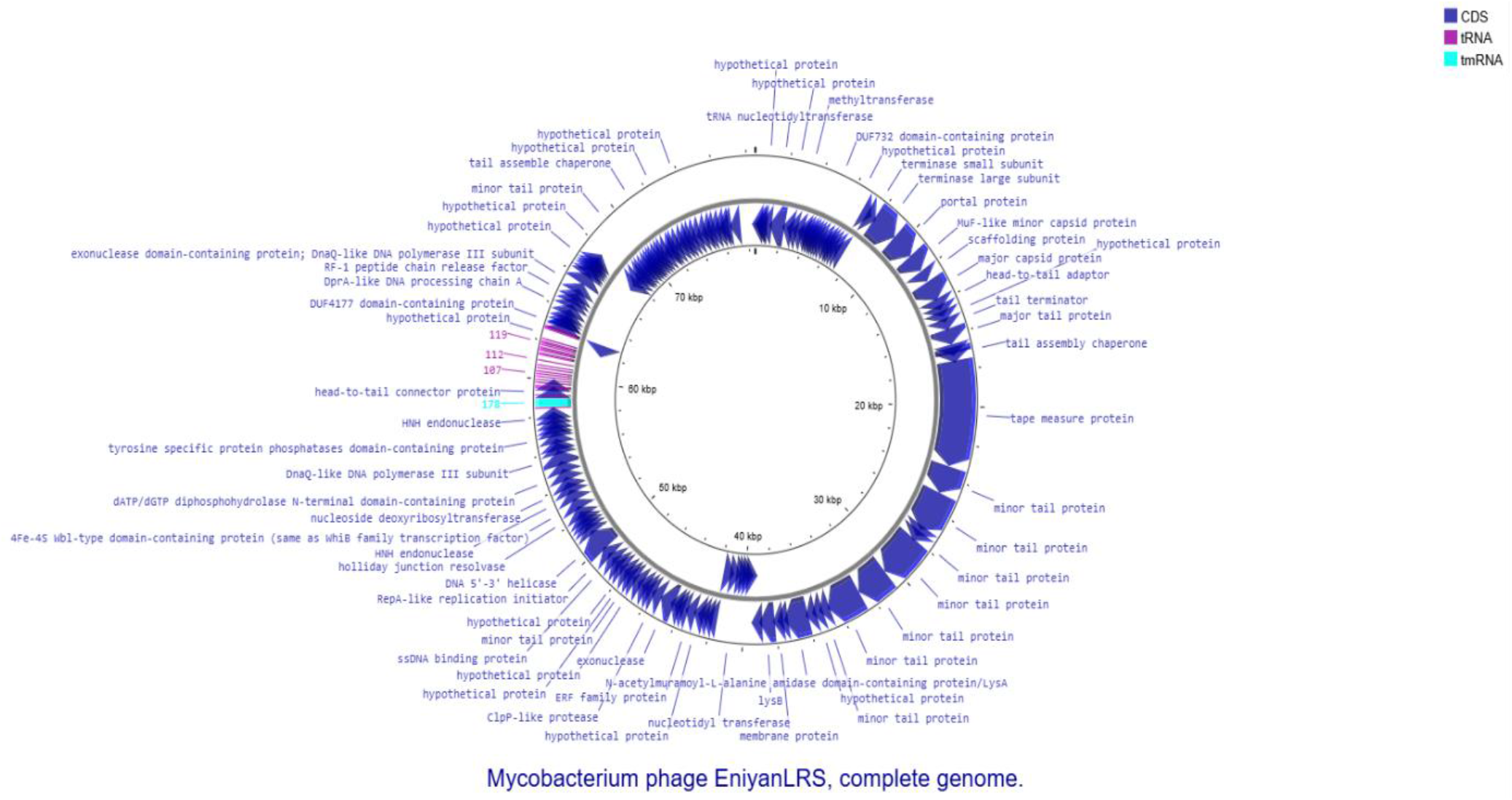
A circular genome map of mycobacteriophage EniyanLRS. The map shows annotated coding sequence (CDS) regions, predicted functions, t-RNA, tm-RNA and GC-skewed regions. **Genome map is generated using the CGView tool.**

**Figure S2.**
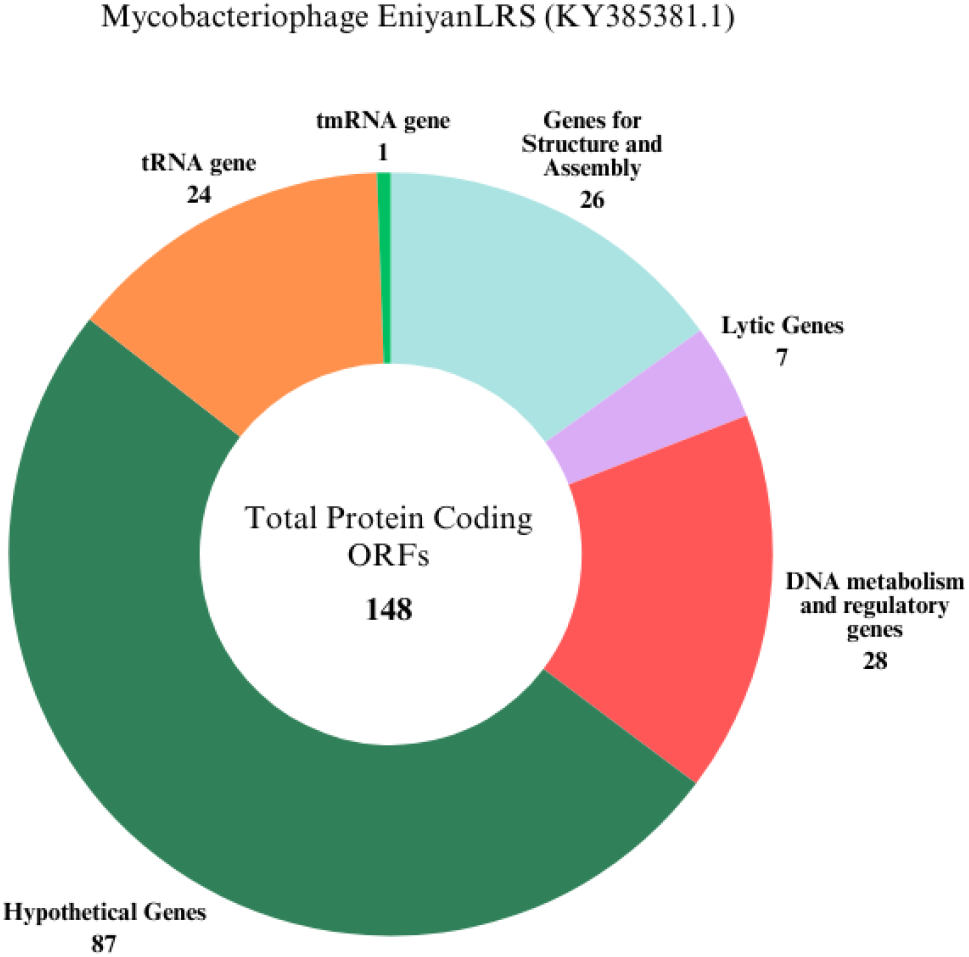
Pie chart representing functional categories in the annotated genome. The genome is categorized based on the predicted functions of ORFs (148) encoding: 1. Structure and Assembly Proteins, 2. Lytic enzymes, 3. DNA metabolism and regulatory proteins, 4. Hypothetical or Uncharacterised proteins, 5. tRNA, 6. tmRNA.

**Figure S3.**
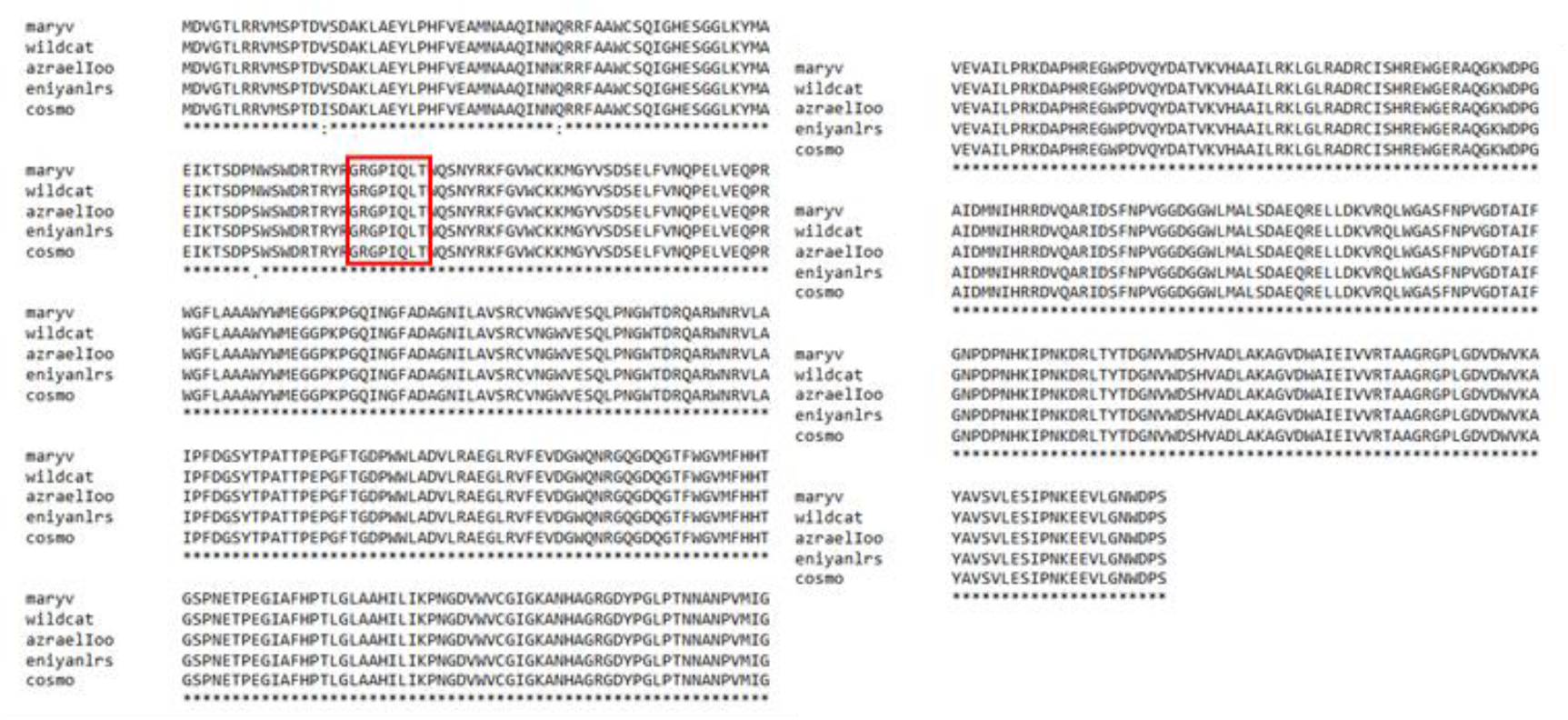
Multiple Sequence Alignment of LysA protein sequences from Cluster V mycobacteriophages. Alignment of LysA sequences from EniyanLRS, MaryV, Wildcat, Azrael100 and Cosmo mycobacteriophages was performed using ClustalW (https://www.genome.jp/tools-bin/clustalw). The conserved G-R-G-P-I-Q-L-T motif, which is associated with chitinase function, is highlighted in a rectangular box. Symbols ‘*’, ‘:’, and ‘.’ denote identical amino acids, conserved substitutions, and semi-conserved substitutions, respectively.

**Figure S4.**
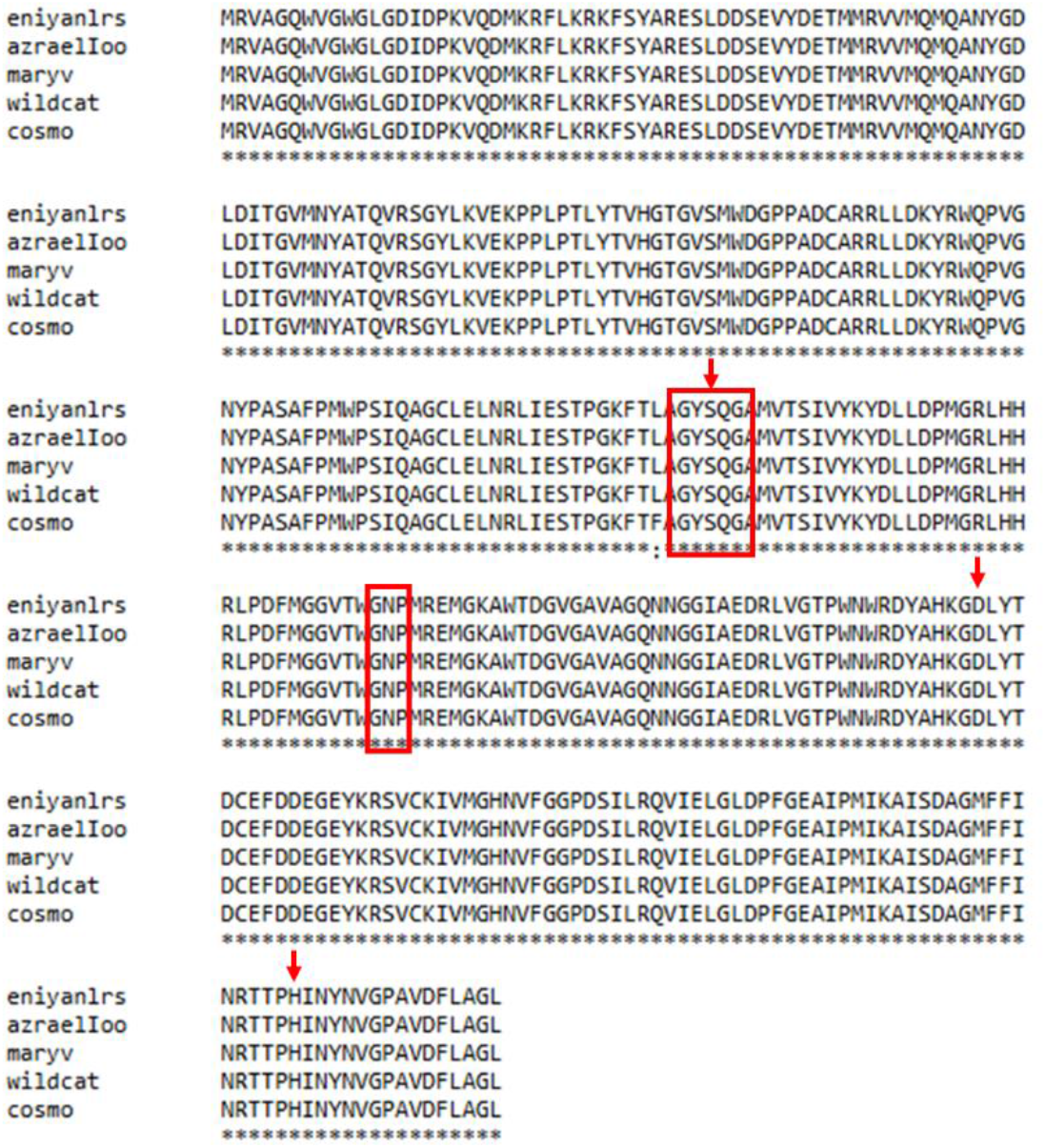
Multiple Sequence Alignment of V LysB protein sequences from Cluster V mycobacteriophages. Sequence Alignment of LysB from EniyanLRS, MaryV, Wildcat, Azrael100 and Cosmo mycobacteriophages was performed using ClustalW (https://www.genome.jp/tools-bin/clustalw). The conserved pentapeptide GXSXG (GYSQG), a characteristic of the serine esterase enzymes and the GXP motif (GNP), a characteristic of cutinases, are highlighted with a rectangular box. The conserved catalytic triad Ser-Asp-His is marked with a red arrow. Symbols ‘*’ and ‘:’ denote identical amino acids and conserved substitutions, respectively.

**Figure S5.**
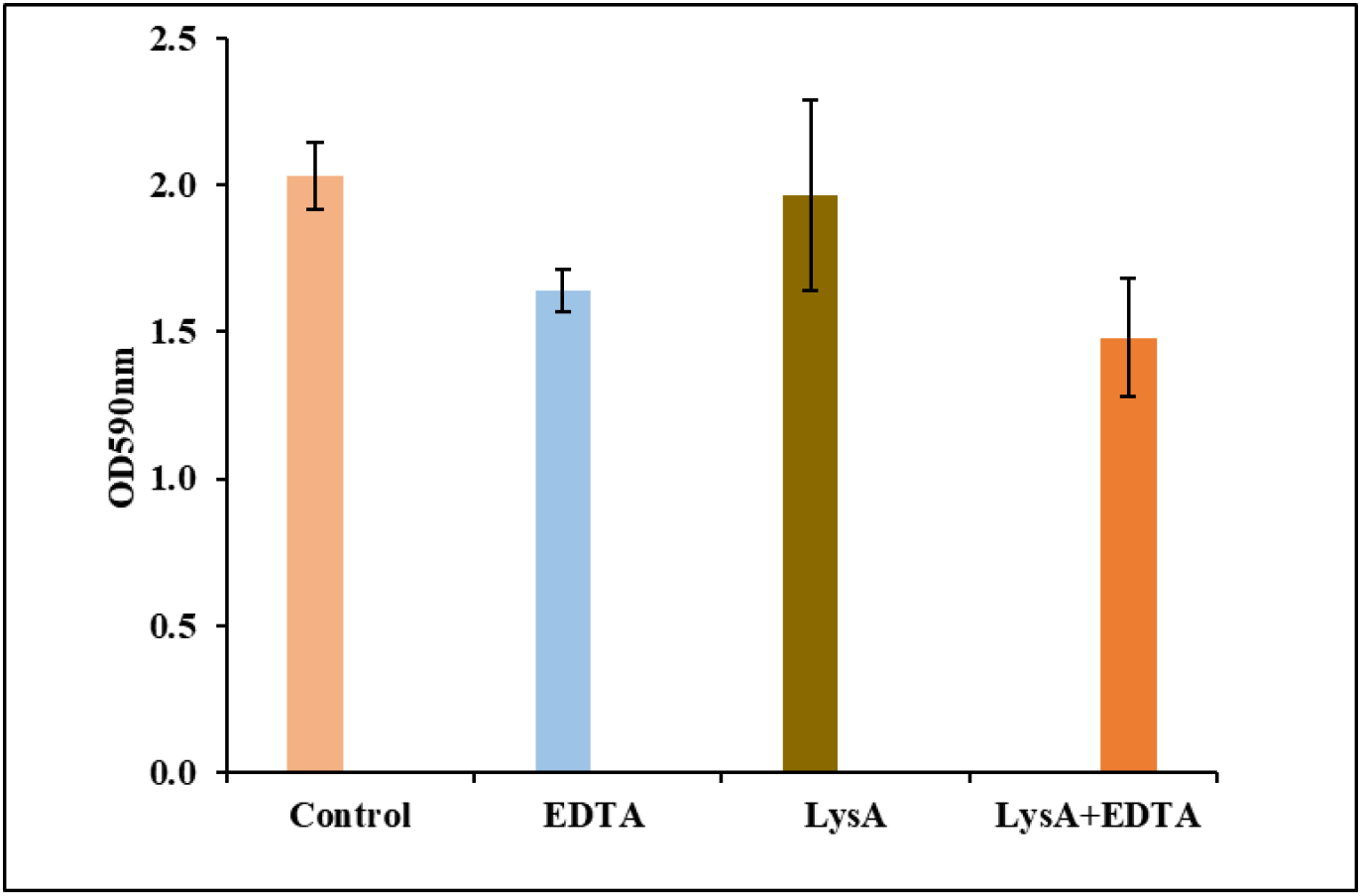
Antibiofilm effect of EniyanLRS LysA in *M. smegmatis* Mc^2^155 using Crystal Violet Assay. The biofilm inhibitory effect of EDTA (3 mM), LysA (2.5 µM) and EDTA + LysA combination (3 mM + 2.5 µM) was measured after 96 h of incubation at 37°C. Controls represent untreated *M. smegmatis* biofilm formation. EDTA, a divalent cation chelator, alone showed 19.4% inhibition; LysA alone did not exhibit biofilm inhibitory effect; EDTA + LysA combination showed 27.17% inhibition. Data is presented as the mean of three independent experiments performed in triplicate (n = 9).

## Data availability

The genome sequence of EniyanLRS is available at GenBank (https://www.ncbi.nlm.nih.gov/genbank/) under the accession number KY385381.1.

## Author contributions

**Kanika Nadar:** Writing – original draft & editing, Methodology, Experiments, Data curation, Visualisation, Validation. **Kandasamy Eniyan:** Writing – review & editing, Methodology, Experiments, Data curation, Validation. **Urmi Bajpai:** Conceptualisation, Funding acquisition, Writing – review & editing, Validation, Supervision, Project administration, Methodology, Investigation.

## AI declaration

The authors employed Google Gemini (OpenAI) for language refinement and manuscript organisation. After using these tools, the authors reviewed and edited the content as needed and take responsibility for the content of the published article.

## Acknowledgements

We thank Anusandhan National Research Foundation (ANRF), previously Science and Engineering Research Board (SERB), GoI, New Delhi, India, for supporting the project (EMR/2017/004051). Kanika Nadar received the Innovation in Science Pursuit for Inspired Research (INSPIRE) fellowship from the Department of Science and Technology (DST), India. Kandasamy Eniyan received the TCOF fellowship from the Council of Scientific and Industrial Research (CSIR). We thank Acharya Narendra Dev College (ANDC), University of Delhi, New Delhi, India, for providing the infrastructural support. We express our sincere gratitude to Prof Graham F. Hatfull and Dr Jacobs-Sera Deborah, Department of Biological Sciences, University of Pittsburgh, USA, for their insightful suggestions and comments that contributed to the preparation of the manuscript.

